# Phyllosphere microbiome-based biocontrol solution against *Botrytis cinerea* and *Plasmopara viticola* in grapevine

**DOI:** 10.1101/2025.02.10.637449

**Authors:** Nicolas-David Rappo, Sylvain Schnée, Clara Chevalley, Émilie Michellod, Floriane L’Haridon, Katia Gindro, Laure Weisskopf, Sébastien Bruisson

**Author notes:** Funding: co-financed by Innosuisse; co-financed by the Swiss National Science Foundation (grant 310030 207917 to LW). Corresponding author: S. Bruisson; Chemin du musée 10, CH-1700 Fribourg, Switzerland.

## Abstract

Plants harbor different microbial communities in their different organs. This difference is due to the various conditions to which the different parts of the plant are subjected. Consequently, the phyllosphere microbiota can differ strongly from root-associated communities. We hypothesize that the grapevine phyllosphere is a valuable source of biocontrol bacteria, whose adaptation to the aerial plant environment may enable them to reach their full protective potential against foliar pathogens. In previous work, we isolated phyllosphere bacteria and showed that many strains were very effective against such pathogens *in vitro*, inhibiting their mycelial and spore development. This work investigates the biocontrol ability of these phyllosphere bacteria in leaf disc assays against two foliar pathogens: *Botrytis cinerea* (gray mold) and *Plasmopara viticola* (downy mildew). Our results show that 40 strains out of 46 affected at least one pathogen by altering their spore physiology and/or reducing disease progression. Among these strains, 27 strains could impact both pathogens and 20 were also capable of stimulating plant defenses. Because bacterial consortia might perform better than single strains, we also compared the protection conferred by individual bacteria and associations of up to three strains. When combined, bacteria showed improved efficacy *in vitro* against *B. cinerea* and *in planta* against *P. viticola*, resulting in a much stronger protection than that obtained by the application of the strains alone. These data suggest that phyllosphere bacteria could be a promising tool to help winegrowers manage grapevine diseases by providing an effective and sustainable protection of their crops.

## Introduction

Plant microbiota forms a rich and complex network that is organized into different communities. Each part of the plant provides a specific environment with conditions favorable to the development of different species of bacteria (Compant et al., 2019). Consequently, within the same plant, bacterial communities found in roots are different from those found in leaves and flowers (Zarraonaindia et al., 2015). Furthermore, because the composition of each community is determined by the sum of all conditions they face, the bacteria found at the surface or inside the plant of the same organ are also different (Dastogeer et al., 2020). Many recent studies have shown that the plant microbiota contributes to its host’s health by modulating its growth, promoting plant defenses and inhibiting pathogen development (Brader et al., 2017; Rai et al., 2023) and such bacteria are good candidates for developing biocontrol solutions.

However, among the thirty to forty current biological control solutions used in Europe and the USA against crop diseases, most rely exclusively on organisms isolated from the soil or roots, even when the objective is to control pathogens affecting the aerial parts of the plant. Of all the solutions available, relatively few organisms have been isolated from the aerial parts of plants such as the yeast *Aureobasidium pullulans* (Botector® and BlossomProtect™), the yeast *Metschnikowia fructicola* (Noli), and the bacterium *Pantoea agglomerans* (Bloomtime Biological™) (Borges et al., 2021; Pascale et al., 2020; Syed Ab Rahman et al., 2018).

In this study, we tested the potential of bacteria isolated from the phyllosphere (the leaves and their surrounding environment) to protect plants against diseases affecting their aerial parts. We selected grapevine as a model plant to test this strategy, for several reasons: i) it is a crop of major economic importance worldwide; ii) it is a perennial plant, and such plants are expected to harbor more adapted phyllosphere microbiota compared to annual plants, since perennial plants have longer-lasting interactions with their associated bacterial communities (Primieri et al., 2022).In such plants, even though leaves are deciduous, the leaf microbiota can be renewed each year from persistent reservoirs present on the plant such as bark (Griggs et al., 2021; Martins et al., 2013; Vitulo et al., 2019); iii) most grapevine cultivars are highly susceptible to leaf-borne pathogens, making this crop one the most fungicide-intensive in the world (W. J. Li et al., 2011). Biocontrol is of particular interest for such cultures that need alternative and novel effective solutions, to limit the environmental toxicity linked to the excessive use of phytosanitary products. In addition, the use of microorganisms that are antagonistic to filamentous plant pathogens is based on multiple modes of action. It is therefore unlikely to lead to the emergence of resistant individuals within pathogen populations and represents a sustainable means of control.

Biocontrol bacteria have a wide range of modes of action and can act either directly on the pathogens and/or indirectly, through the plant. Therefore, they can have a direct inhibitory effect through the production of metabolites that can disturb the pathogen’s growth and physiology, impair their virulence, or they can also compete for space and nutrients, e.g. by trapping vital resources such as iron through the secretion of siderophores (H. Chen et al., 2008; Haas & Défago, 2005; Ongena & Jacques, 2008). They can also have an indirect effect on the pathogen by improving plant health through the production of plant growth promoting compounds or the induction of a local or systemic defense response (Legein et al., 2020). In grapevine, one of the well-identified metabolic defense pathways is based on the production of stilbenic compounds, a group of phytoalexins of major importance for this crop due to their high toxicity for fungal pathogens (Pezet et al., 2003, 2004). This family includes a large number of inducible molecules that are produced in several forms from monomers to tetramers (Schnee et al., 2013). Their distribution and concentration are cultivar and organ dependent and the resistant cultivars have been shown to synthesize high amounts of the more active forms such as viniferins (Waffo-Teguo et al., 2008; Viret et al., 2018; Guerrero et al., 2020).

In view of these multiple modes of action, combining different strains with complementary bioactivities into a mixed microbe-based biocontrol solution could provide a multi-layer protection by combining several mechanisms that may lead to improved efficacy and increased plant protection. Several studies already highlighted that consortia of beneficial microbes can help to improve the effectiveness of the protection against pathogens in different cultures such as strawberries and tomatoes (Guetsky et al., 2001; Maciag et al., 2023; Niu et al., 2020). Additionally, the use of consortia can help to make the protection less variable from one plant to another, compared to the use of single strains (Yu et al., 2022; Wang et al., 2023).

In a previous study, we demonstrated that the grapevine microbiota isolated from the phyllosphere contained many effective strains that could effectively inhibit the development of leaf pathogens in *in vitro* experiments (Bruisson et al., 2019). In this previous work, several bacteria were observed to strongly inhibit the mycelial growth of the fungus *B. cinerea* and the oomycete *Phytophthora infestans* in *in vitro* conditions. They were also able to disrupt the physiology of these pathogens by affecting their spores. We concluded that the use of such bacteria was a promising strategy to increase the effectiveness of microbe-based biocontrol solutions, but that more advanced testing using whole plants was required.

In the present work, we aimed to determine whether grapevine phyllosphere bacteria can confer *in planta* protection and whether the application of phyllosphere bacteria in consortia provide, as hypothesized, superior protection through the addition of individual effects compared to the use of single strains. We tested these hypotheses against two of the most detrimental grapevine diseases: downy mildew caused by the oomycete *Plasmopara viticola*, and gray mold caused by the ascomycete *Botrytis cinerea*. Our results suggest that this protection is achieved by combining direct effects on the pathogens, affecting their spores and their mycelium development, as well as through indirect effects linked to their ability to increase the expression of defense-related genes and the production of specialized metabolites involved in plant defense.

## Materials and methods

### Biological material and culture media

Bacterial strains isolated from grapevine phyllosphere as described in (Vionnet et al., 2018) and screened for their biocontrol capacity in (Bruisson et al., 2019) were grown on LB (Lysogeny Broth) medium in the dark and at room temperature. LB medium was prepared by mixing 12.5 g/L of LB Broth Miller and 10 g/L of LB Broth Lennox (Fisher Bioreagents) in distilled water. For solid medium, 15 g/L of Agar-Agar Kobe I (Roth) were added.

All bacteria are epiphytic or endophytic strains isolated from three different grapevine cultivars. The first two letters of their code name correspond to their cultivar of origin: Chasselas (CH), Pinot Noir (PI) and Solaris (SO). The last letter corresponds to their location of origin: D for endophytic and P for epiphytic.

*Botrytis cinerea* strain 2905 was provided by the Agroscope of Changins and was grown in potato dextrose agar (PDA) (Roth). PDA was prepared by dissolving 39 g/L of PDA powder (Sigma-Aldrich) in distilled water.

*Plasmopara viticola* sporangia were provided by the Agroscope of Changins. They were harvested from naturally infected leaves collected yearly in local vineyards to reflect the genotypes currently infecting the grapevine at the time of the experiments. These field isolates were maintained by reinoculation on Cabernet Sauvignon.

The plant material studied was based on cuttings of *Vitis vinifera* L. cv. Gamay (susceptible to *B. cinerea*) and Gamaret (susceptible to *P. viticola*). After 2 months of growth in greenhouse conditions, plants were transferred into growth chambers for infection assay, with a 16 h/8 h photoperiod, 21/20°C day/night temperature and 40 % relative humidity (RH). A sulfur lamp was provided to prevent the appearance of powdery mildew.

### Effect of bacterial strains on pathogens

Experiments to assess the effects of grapevine phyllosphere bacteria on *B. cinerea* and *P. viticola* can be divided into three main parts (Fig. 1). The first assays consisted in testing the ability of the bacterial strains to disrupt the pathogens’ reproduction by interfering with their spore development. Co-incubation with *B. cinerea* conidia and *P. viticola* zoospores was used to monitor the strains’ ability to inhibit their germination and motility, respectively. After co-incubation with the bacteria, *P. viticola* zoospores were used to infect leaf discs to evaluate if the loss of motility impaired their infectivity. In the second assays, the protective abilities of the bacteria were tested by application of the strains on leaf discs two days prior infection in order to let them adapt to their new environment and the possibility of stimulating plant defenses, before measuring their protective effect. After a selection step based on the effectiveness of the strains in both series of assays, their resistance to biofungicides and their compatibility, a last assay on whole plants was conducted. This was a preventive assay to test the local and systemic protection conferred by a selection of bacteria used alone or in combinations of three strains to evaluate their protective abilities.

**Figure 1:**
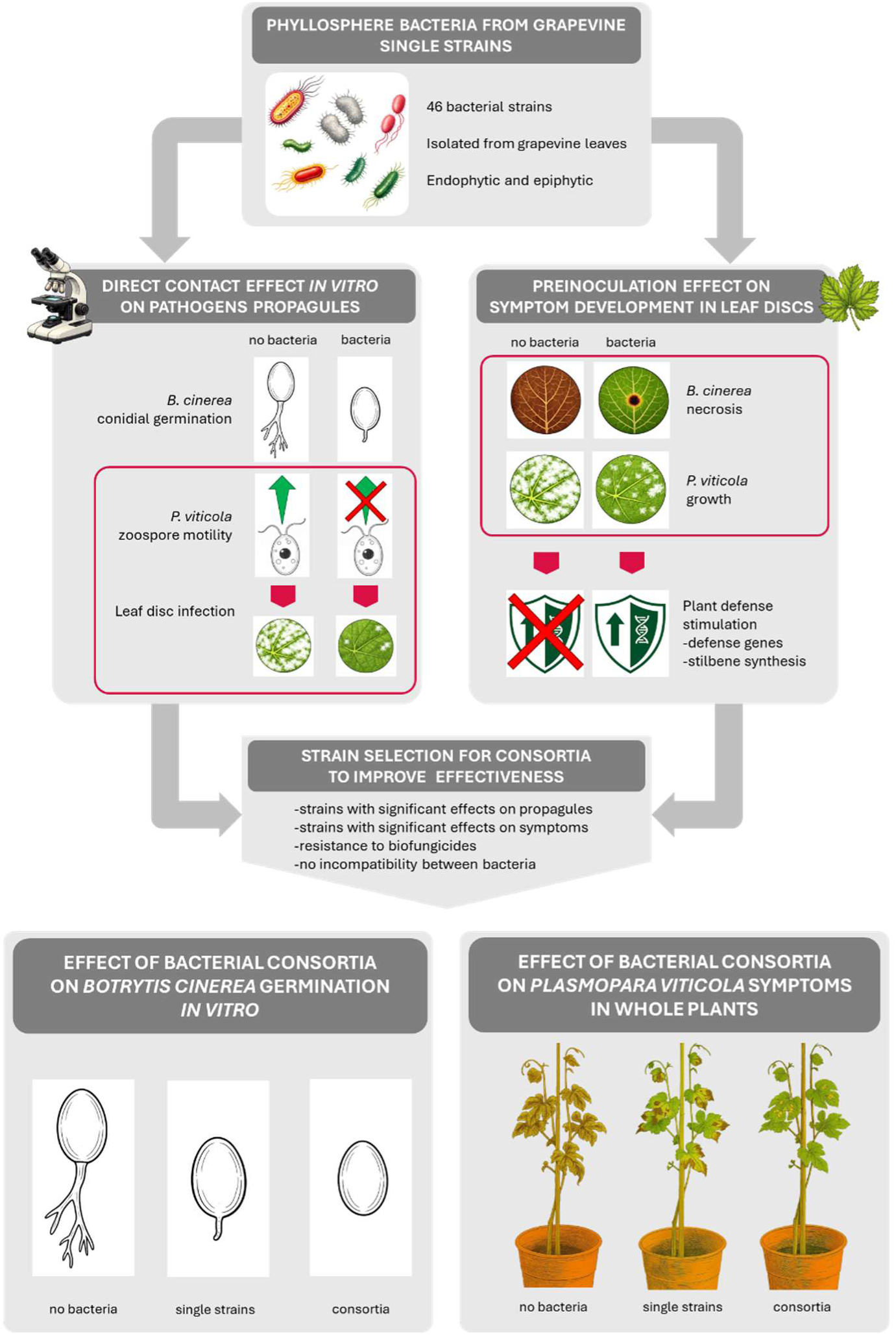
Summary of the experimental workflow: *in vitro* experiments against pathogen propagules, leaf disc assays, selection of bacteria for consortia and consortia effect on propagules and whole-plant infections. Red boxes indicate experiments that are part of the same experimental setup; a red cross over an icon indicates absence of the respective effect. Illustration developed with assistance from Sora (OpenAI), with final graphics produced by the authors.

### Effect of bacterial strains on *B. cinerea* spore germination

Spores of *B. cinerea* from 7-day-old mycelium grown on PDA plate were harvested by watering the plate with 5 mL of sterile water and filtration through glass wool to remove hyphae. The spore solution was then adjusted to 5 x 10^4^ spores/mL and 12 µL of this solution was mixed in an Eppendorf tube with 36 µL of bacterial suspension adjusted at OD_600_ = 1 in V8 medium that was filtered using a 0.2 µm Millex®-FG filter (Merck). Each single strain from our collection was tested at OD_600_ = 1. Combinations of two to three strains were tested using equal volumes of bacterial solutions with a final concentration of OD_600_ = 1.

The mixture was transferred into a 24-well plate, and the plates were incubated at 21°C under light for 8 hours. Three replicates were performed for each modality and the control consisted of spore solutions mixed with filtered V8 without bacteria. After 8 h pictures were taken using an imaging plate reader (Cytation5, Biotek). Spores were counted according to their development stage (not germinated, normal germination, delayed germination). Each assay was replicated twice with n = 3 repetitions per experiment.

### Effect of bacterial strains on *P. viticola* zoospore motility

Zoospores of *P. viticola* were released from sporangia harvested by suction from sporulating lesions of artificially inoculated leaves (*V. vinifera* cv. Cabernet Sauvignon) with a filter-containing pipette tip connected to a vacuum device. The sporangia were resuspended in sterile water and their concentration was adjusted to 5 x 10^4^ sporangia/mL in a falcon tube. The tube was placed in a shaker at 25 rpm at room temperature and in the dark for zoospore release. After 1 h, the zoospore solution concentration was adjusted to 5 x 10^4^ zoospores/mL. The effect of bacteria on zoospore motility was assessed by mixing 500 µL of a solution of bacteria diluted at OD_600_ = 2 in sterile water with 500 µL of the zoospore solution in a 1.5 mL tube for a final concentration of OD_600_ = 1 (1:1 mixing). The solution was then incubated at room temperature in a shaker at 25 rpm for 30 minutes. After incubation, the solution was transferred into a counting chamber and observed under a microscope to visually count the number of motile and non-motile zoospores. After counting, the percentages of motile and non-motile zoospores were calculated and compared to the negative control (zoospore mixed with distilled water only) and the positive control (zoospore mixed with a 2 g/L solution of copper hydroxide (Kocide® Opti, Bayer). For each strain, two independent experiments were performed with n = 3 replicates per experiment and each replicate containing at least 50 zoospores. In order to determine the effect of the loss of motility on the zoospore’s infectivity an aliquot of the suspension combining zoospores and individual bacteria was then used for leaf disc infection after the 30-minute incubation, following the procedure described below in the leaf disc infection section.

### Electron microscope observation of zoospores

Pellets of the different modalities (*P. viticola* zoospores, the bacterial strain PID6 (*Bacillus* cereus), and *P. viticola* zoospores in contact with PID6) from the experiment described above were prepared according to Roland and Vian (1991). Briefly, the pellets were centrifuged for 5 minutes at 2000 rpm and resuspended in a solution of 3 % glutaraldehyde–2 % paraformaldehyde in 0.14 M PIPES buffer (pH 7) for prefixation. Samples were then centrifuged for 20 minutes at 2000 rpm, the supernatant was removed, and the samples were washed twice with 0.07 M PIPES buffer. They were embedded in 2 % agarose and postfixed with a solution of 1 % OsO4. The samples were then dehydrated in a graded series of ethanol solutions (30–50–70–95–100 % v/v) and infiltrated in a solution of ethanol mixed with propylene oxide (1:1) and pure propylene oxide, followed by a series of epoxy resin mixed with propylene oxide (1:3, 1:1) for 2 hours each. Finally, they were infiltrated with epoxy resin for 17 hours. After polymerization for 48 hours at 60°C, a staining step was performed by floating on a 50°C solution of 1 % methylene blue, 1 % sodium borate, and 1 % azure II for 10 seconds for semi-thin sections. Thin sections were cut and stained with 2 % uranyl acetate followed by lead citrate, according to Reynolds (1963). The thin sections were observed with a Tecnai G2 Spirit BioTWIN transmission electron microscope (TEM) equipped with a Gatan UltraScan High-resolution CCD camera.

### Evaluation of the preventive effect of the application of bacteria on gray mold and downy mildew infection on leaf discs

Leaf discs from *V. vinifera* L. cv. Gamay and cv. Gamaret for *B. cinerea* and *P. viticola* respectively were cut from 2-month-old grapevine leaves taken from leaf stages 4 to 7. Leaves were taken randomly from at least ten plants and were rinsed with tap water and surface sterilized by immersion in calcium hypochlorite (40 g/L) for 7 minutes followed by rinsing in sterile water prior to cutting. Leaves were then dried with sterile filter paper, used to cut leaf discs using a 1.7 cm diameter punch and placed in sterile water. For each treatment, three Petri dishes containing ten leaf discs each were used, leaf discs were taken randomly and placed on top of a filter paper moistened with sterile water placed inside a 9 cm Petri dish with the abaxial side facing upward and placed in a growth chamber with a 16 h/8 h photoperiod, 21/20°C day/night temperature and 40 % RH. After 24 h each plate was inoculated by spraying 500 µL of bacterial solution diluted in sterile water and adjusted to OD_600_ = 1, using a sterile glass atomizer and placed back in the growth chamber for two days, to let them adapt and develop in their new environment. The leaf discs were then infected with *P. viticola* or *B. cinerea*. For *P. viticola*, the infection was performed by spraying 50 µL of solution adjusted at 5 x 10^4^ zoospores/mL in each plate. For *B. cinerea*, the infection was performed by depositing a 20 µL droplet of a spore solution adjusted to 1 x 10^5^ spores/mL in PDB^1/4^ (quarter-strength potato dextrose broth) on a wound made by a sterile syringe at the center of the leaf disc. The same procedure was used with sterile water only for the negative control. For positive controls, leaf discs were treated with a 2 g/L solution of copper hydroxide (Kocide® Opti, Bayer) against *P. viticola* and a 0.04 % solution of the yeast *Aureobasidium pullulans* (Botector®, Andermatt) against *B. cinerea*. Petri dishes were sealed with Parafilm® and placed back in the growth chamber. For each modality, one plate on the three available containing 10 leaf discs was used to collect material for gene expression analysis by qPCR and stilbene synthesis by UPLC. Five leaf discs were collected, dried and snap frozen in liquid nitrogen 6 h after infection for qPCR analysis to monitor early changes in defense-related genes expression, and the other five 3 days after infection for UPLC analysis, 3 days being the time required to observe a representative level of stilbene synthesis. Pictures of each remaining plate were taken one week after infection to assess the development of symptoms. Two independent experiments were performed for each modality.

### Whole plant infection with *P. viticola*

Half of the leaves from foliar stage 4 to 7 of two-month-old grapevine plants (4 plants per treatment) were sprayed using a sterile glass atomizer with a solution of 4 mL of bacteria adjusted to OD_600_ = 1 on their abaxial face, while the other half was protected with plastic bags to prevent unwanted inoculation. Once the inoculation was complete, plastic bags were removed and uninoculated leaves were sprayed with sterile water. Two days after the bacterial inoculation, all leaves from foliar stage 4 to 7 were sprayed with 200 µL of a solution of *P. viticola* zoospores adjusted to 5 x 10^4^ zoospores/mL. After a week, leaves were collected, the level of symptoms was quantified and compared to the control (plants inoculated with water only) in search of local and systemic protection from the bacteria. Two independent experiments were performed for each modality.

### Symptom assessment

For symptom measurement on plant tissue, pictures of leaf discs or whole leaves were analyzed in an automated and quantitative manner. A macroinstruction was developed in the freeware ImageJ to measure the necrosis development of *B. cinerea* and another one to measure the formation of sporangiophores of *P. viticola* as described in Guyer et al. (2015). The macro instruction analyzes each leaf tissue in an image. It counts the total number of pixels occupied by the leaf as well as the number of pixels occupied by *B. cinerea* necrosis or *P. viticola* sporangiophores within each leaf tissue. A report is then generated to calculate the percentage of surface area occupied by the pathogen on the surface of each leaf tissue. The level of inhibition was calculated as a percentage of the level of infection observed in the negative control (inoculated with water instead of bacteria). For positive controls, plants were treated with a 0.4 % solution of brown algae (Alginure®, Andermatt) against *P. viticola* to compare the effectiveness of our bacterial treatments against a certified biocontrol solution.

### Gene expression analysis

The ability of phyllosphere bacteria to promote defense gene expression on grapevine was analyzed by qPCR in 8-week-old plants. Five leaf discs from the preventive inoculation assays on leaf discs against *B. cinerea* and *P. viticola* described above were sampled 6 h after infection, snap-frozen in liquid nitrogen and stored at −80°C. RNA was extracted with the NucleoSpin RNA Plant from Macherey-Nagel with a modified procedure (Schellenbaum et al., 2008). Reverse transcription was performed using the SensiFAST cDNA Synthesis Kit from Bioline. Quantitative PCR reactions were performed using the SensiFAST SYBR Hi-ROX Kit from Bioline. Each reaction was carried out using a Mic qPCR Cycler (bio molecular systems) and the software micPCR v2.8.13 with 5 µL of cDNA (5 ng/µL), 7.5 µL of SYBR Hi-ROX mix, 0.5 µL of each primer and 1.5 µL of sterile water. For amplification, an initial denaturation step at 95°C for 15 min was done, followed by 45 amplification cycles (95°C for 15 s, 60°C for 15 s, 72°C for 30 s). The experiment was repeated twice, with n = 3 samples in each experiment. For each reaction a technical replicate was performed and values with a standard deviation lower than 0.25 (or 0.5 Cq) were averaged and used for further calculations. All results were analyzed using the double delta Cq method with infected samples treated with water as reference and normalization was performed relative to the expression of the two reference genes Actin and 60SRP (60S ribosomal subunit). After normalization and calculation of the fold change, results were log2 transformed to make fold-changes symmetric.

Primers were used to monitor the expression of the following defense-related genes, PR-1 (pathogenesis-related protein 1), PR-4 (pathogenesis-related protein 4), STS (stilbene synthase), ROMT (resveratrol O-methyltransferase), PAL (phenylalanine ammonia-lyase) and LOX (lipoxygenase) (Table S1).

### Stilbenic compound measurement

Leaf discs were cut in small pieces with a scalpel, weighed and placed into microcentrifuge tubes. Samples were immediately snap-frozen and immersed in 300 μL of methanol. The tubes were then heated at 60°C for 15 minutes with agitation at 800 rpm, followed by rapid cooling in an ice bath for 5 minutes. After cooling, the samples were centrifuged to separate the supernatant. The methanolic extracts were analyzed using Ultra-Performance Liquid Chromatography (UPLC) to quantify the presence of various stilbenic compounds, including trans-piceid, trans-resveratrol, α-viniferin, δ-viniferin, ε-viniferin, isohopeaphenol, and pterostilbene, following the method described by Pezet et al. (2003). The analysis was performed using a Vanquish UPLC system (Thermo Scientific) with a mobile phase consisting of H_2_O with 0.1 % formic acid (Phase A) and acetonitrile with 0.1 % formic acid (Phase B). The gradient elution program started with 5 % B and increased to 40 % over 17 minutes, followed by an increase to 50 % B between 17 and 20 minutes. From 20 to 25 minutes, the proportion of B increased to 100 %, which was maintained until 29 minutes. The flow rate was set at 0.3 mL/min. Separation was carried out using a Hypersil Gold column (1.9 μm, 100 x 2.1 mm, Thermo Scientific) equipped with a 0.2 μm UPLC filter (Thermo Scientific), maintained at 30°C. The injection volume was 2 μL, and detection was performed using a UV detector at 280 nm and 310 nm. This method ensures accurate and reproducible quantification of stilbenic compounds in leaf tissues.

### Bacterial survival in presence of biocontrol products

For the consortia selection process, the ability of the strains to survive in presence of Armicarb® (potassium bicarbonate, Andermatt) and Thiovit® Jet (wettable sulfur, Syngenta), two biocontrol compounds commonly used in viticulture, was assessed. Bacteria were grown overnight in liquid LB at 28 °C and the solution then adjusted to OD_600_ = 1 and used as inoculum for liquid LB medium containing recommended working concentrations of Armicarb (0.2 %), Thiovit Jet (0.4 %) or a combination of both to test their effects on bacterial growth. After 24 hours, the growth of bacteria in contact with the biocontrol products was compared with the growth of the controls (bacteria grown in medium without biocontrol products). Two independent experiments were conducted for each assay.

### Bacterial compatibility assay

Bacteria compatibility was tested in liquid coculture. Each strain was grown overnight in liquid LB at 28 °C and the solution was adjusted to OD_600_ = 1. Then, 50 µL of each strain was used for a coculture in a final volume of 200 µL of LB in a 96-well plate and incubated for 24 hours at 28 °C. The mixture was plated on LB agar supplemented with selective antibiotics (kanamycin, cefotaxime, and gentamicin) chosen based on each strain’s natural antibiotic resistance. Bacteria were considered compatible when each strain was able to grow successfully in selective media after coculture.

### Data analysis

All statistical analyses were performed in GraphPad Prism 10 (GraphPad Software, San Diego, CA, USA). Spore germination assay results (number of spores for each category and for each strain) were analyzed using a Fisher’s exact test with Bonferroni’s correction for multiple comparisons. For motility assays, the differences in the number of motile zoospores treated with bacteria were assessed by using a One-way ANOVA followed by a Dunnett’s test with negative controls (zoospores treated with water) as a reference. For symptom analysis, the effect of the bacteria on symptom development was assessed with an ANOVA on the area of the symptomatic tissue followed by a Dunnett’s test with negative controls (leaf discs treated with water) as a reference.

For gene expression analysis, an ANOVA analyzing the effects of the presence of bacteria in infected plants followed by a Dunnett’s test with negative controls (leaf discs treated with water and infected by pathogen) as a reference, was performed for statistical analysis. Results of all statistical tests are displayed as follows: - = not significant; ∗ = *p* < 0.05; ∗∗ = *p* < 0.01 and ∗∗∗ = *p* < 0.001.

## Results

### 1. *In vitro* assays/Bacterial effects on pathogen propagules

The ability of phyllosphere bacteria to disrupt the germination of *B. cinerea* spores was assessed by confrontation tests. After being in contact with the bacteria for 8 h, the spores displayed four different morphologies: i) normal germination, with expected formation of a long germ tube, ii) partially inhibited germination, with only the tip of the germ tube visible, iii) complete inhibition of germination, with full inability for the spore to form a germ tube, iv) in a few rare cases, spores could also form a swollen germ tube, revealing an abnormal germination in which the germ tip swelled, stopping its elongation (Table 1, Fig. 2 B-E).

**Table 1:**
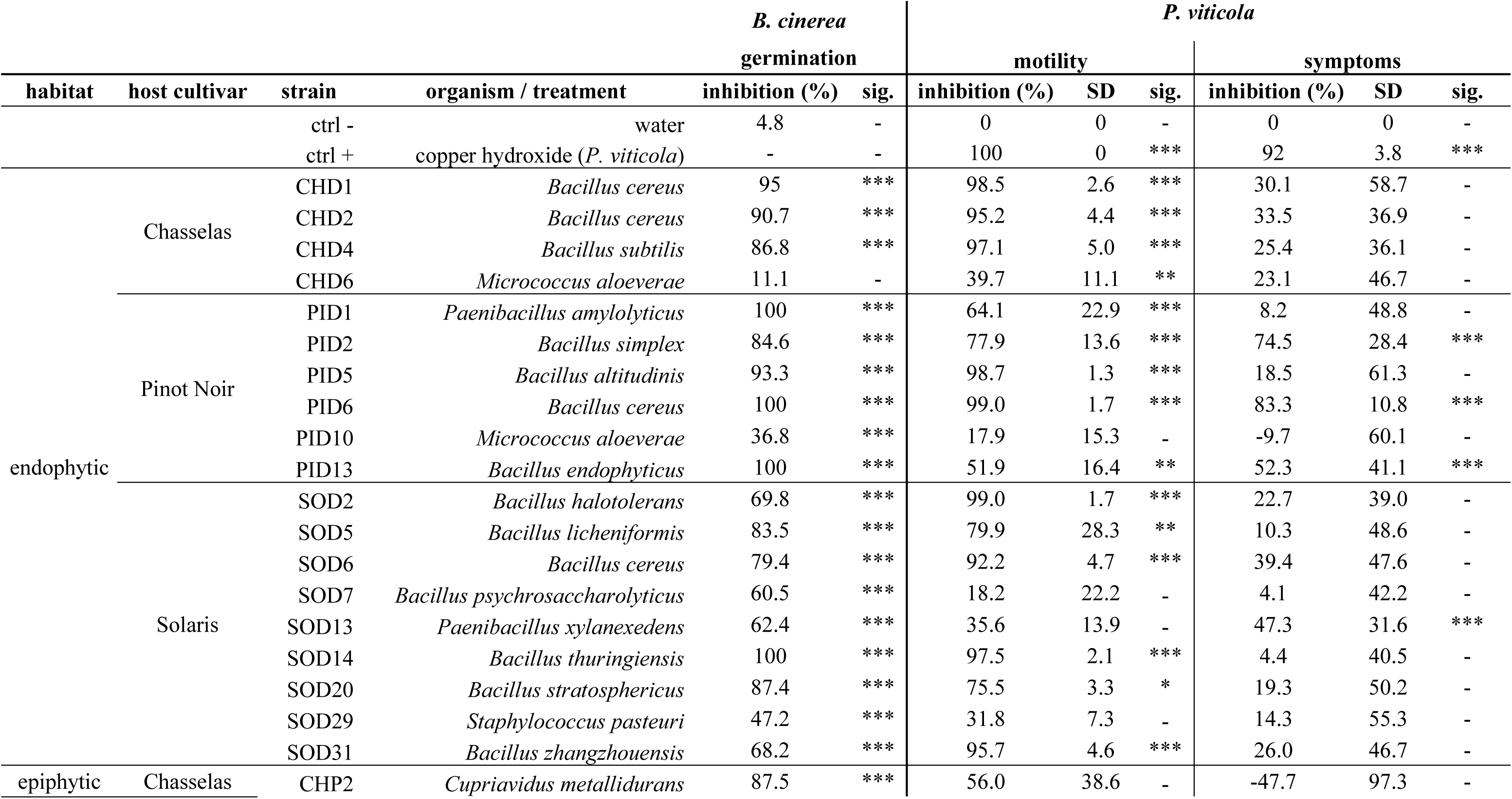
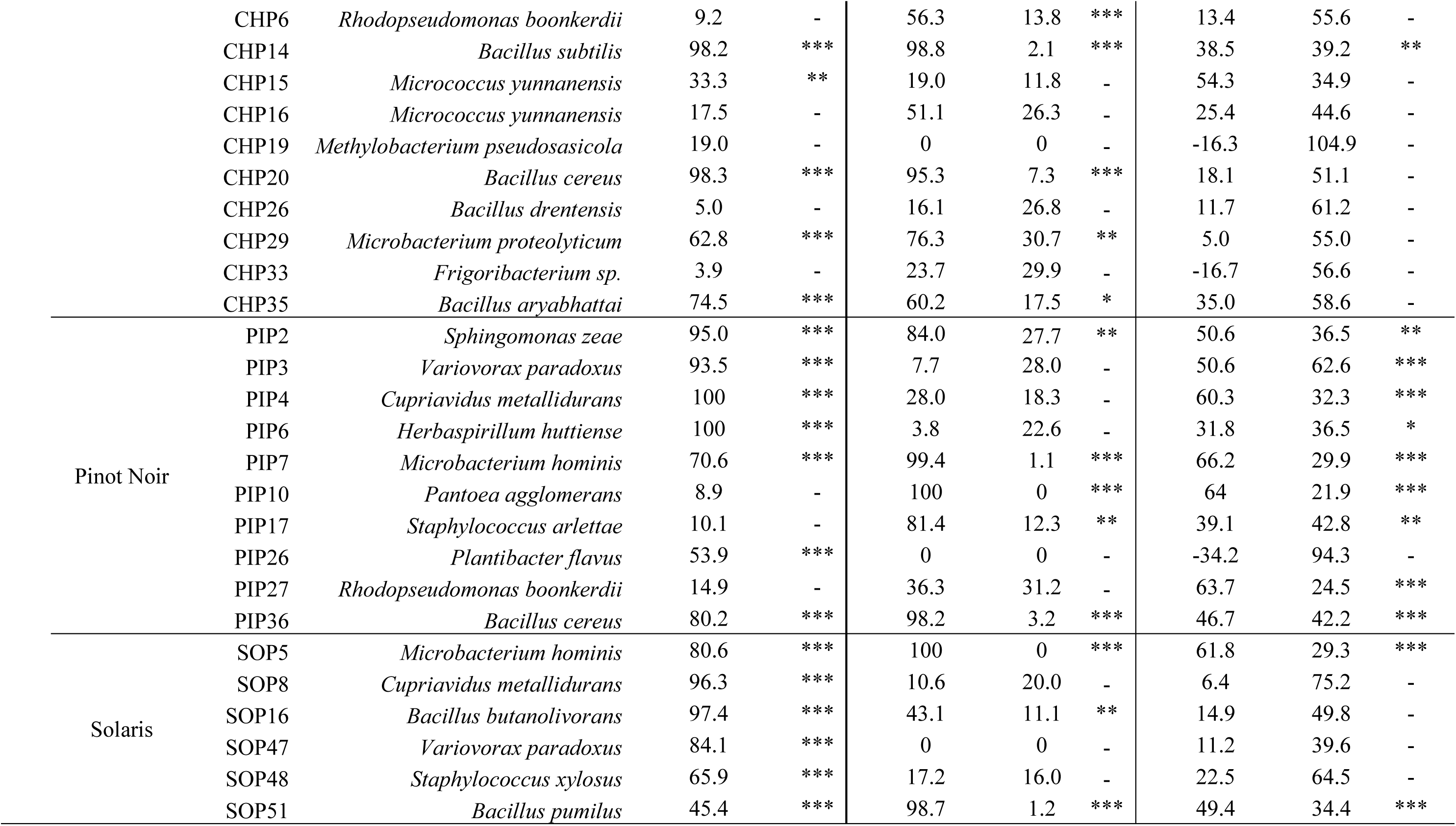
Effect of grapevine-associated phyllosphere bacteria on propagules of *B. cinerea* and *P. viticola*. *B. cinerea* conidial germination inhibition is expressed as the percentage of spores with impaired germination. Activity against *P. viticola* zoospores is reported as the percentage of non-motile zoospores and as the inhibition of symptom development on grapevine leaf discs using the same zoospore suspension exposed to bacteria as in the motility assay. Statistical significance: * = p < 0.05; ** = p < 0.01; *** = p < 0.001; - not significant.

**Figure 2:**
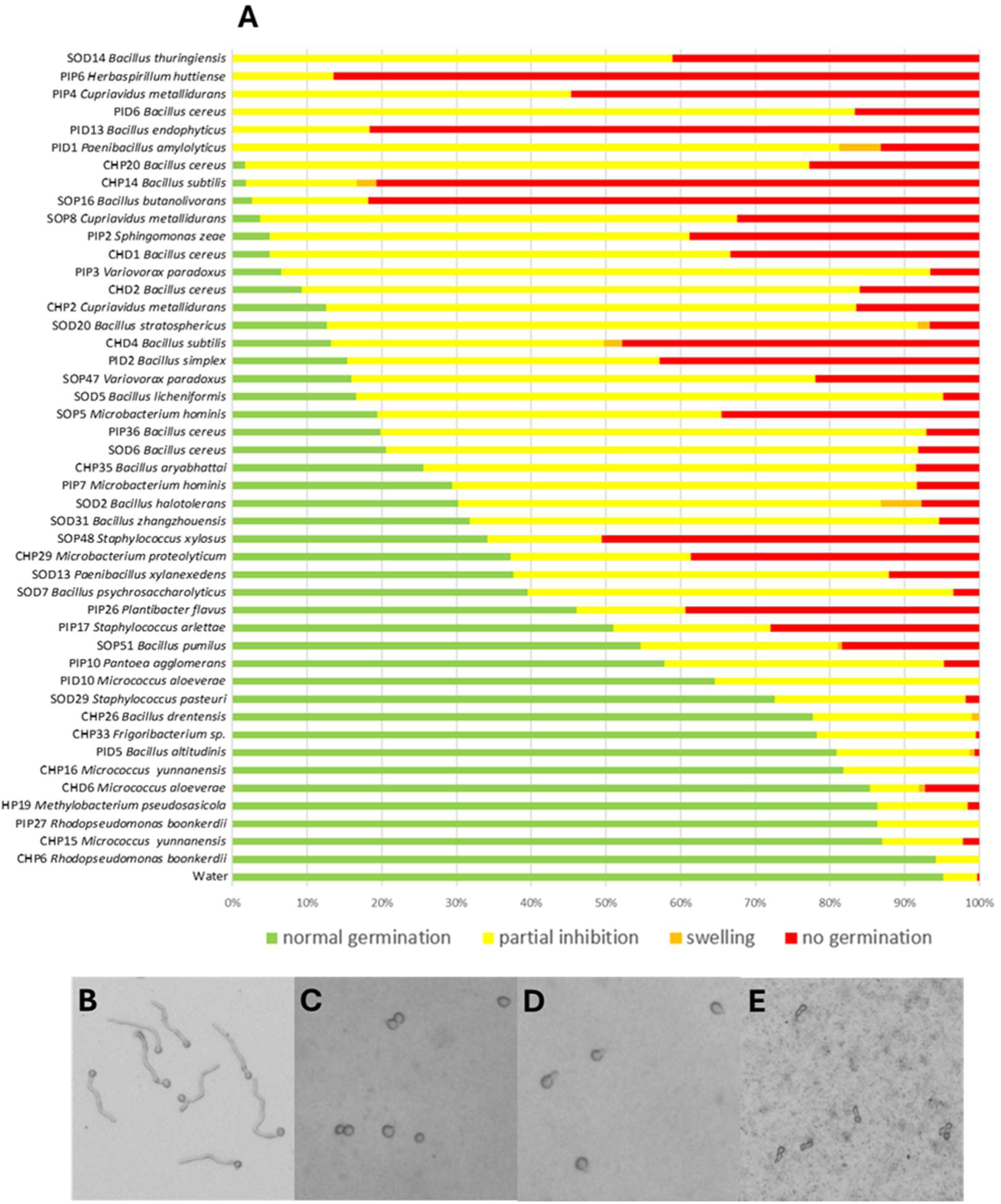
Effect of phyllosphere bacteria on *B. cinerea* spore germination. A: Each colored bar represents the proportion of spores showing the corresponding morphology. Green, normal germination; yellow, partial inhibition (only the tip of the germ tube is visible); orange, swelling; and red, no germination. The different types of germination morphologies are shown in (B to E); B: normal germination; C: absence of germination; D: partial germination and E: swelling.

Among the 46 bacteria that were tested, 33 strains were able to strongly disturb the normal germination of more than 50 % of *B. cinerea* spores, either by strongly delaying the formation of the germ tube or by fully preventing spore germination (Fig. 2 A). The six most effective strains SOD14, PIP6, PIP4, PID6, PID13 and PID1 were even capable of inhibiting the germination of 100 % of the spores, leaving no spore able to germinate properly. These strains belonged mainly to the *Bacillus* or *Paenibacillus* genus, with the exception of PIP4 (*Cupriavidus*) and of the most effective one, PIP6 (*Herbaspirillum*). Five of the six most effective strains were isolated from Pinot Noir.

All bacterial strains were tested to assess their ability to disturb *P. viticola* zoospores *in vitro*. Confrontation assays between bacteria and *P. viticola* zoospores showed that the contact with bacteria could strongly affect zoospore motility. A significant reduction was measured with 28 strains out of the 46 tested. Visual assessment revealed that zoospores either completely lost their motility, or remained motile as in the non-treated controls, but we did not observe slower movement or reduced dispersal. However, the proportion of affected zoospores was different depending on the bacterial strain used. Among these 28 motility inhibiting strains, 16 strains caused a very strong reduction, affecting 90 % to 100 % of the zoospores and reaching the level of inhibition achieved with the copper used as a positive control, while 12 strains caused a significant but lower reduction (40 % to 90 %) (Table 1). A large predominance of bacteria from the *Bacillus* genus were found among the active strains, since 19 of the 21 *Bacillus* strains caused a significant effect. However, the two most effective strains affecting both 100 % of the zoospores (PIP10 and SOP5) were a *Pantoea agglomerans* and a *Microbacterium hominis* strain, respectively. The active strains were composed of a balanced proportion of epiphytic (13) and endophytic (15) strains.

All the suspensions composed of zoospores and individual bacterial strains were thereafter used to infect leaf discs from the susceptible cultivar Gamaret to monitor pathogen development. This was done to test whether the observed loss of motility would reduce the infectivity of *P. viticola* zoospores. The observations showed that in the case of 16 strains out of 46, the infections performed with the treated zoospores resulted in a significant reduction in symptoms (from ca. 30 % to 80 % of the control) compared with the infection performed with non-treated zoospores (Table 1). From these 16 strains, only 11 were also significantly impairing motility, suggesting that the loss of motility was not always correlated with symptom reduction and that other mechanisms are involved. Indeed, five strains that had not impaired zoospore motility were nevertheless effectively reducing the symptoms caused by the zoospores with which they had been co-incubated. Interestingly, four of these five strains are epiphytes, isolated from Pinot Noir and belonged to different genera than those highlighted previously, namely *Rhodopseudomonas*, *Cupriavidus*, *Variovorax* and *Herbaspirillum*.

It seemed that when bacteria (12 strains) isolated from the cultivar Pinot Noir were co-inoculated with zoospores, this almost systematically resulted in significant symptom reduction, whereas strains isolated from the two other cultivars harbored a lesser proportion of protective strains, judging from this leaf disc experiment.

The zoospores that had been in contact with the most effective bacterium PID6 (*B. cereus*) were observed using a transmission electron microscope (TEM). The zoospores that were in contact with non-active bacteria and had not lost their motility displayed a standard oval shape comparable to the control zoospores. However, the zoospores that had been in contact with PID6 showed an irregular shape, suggesting that the zoospore cell had lost its integrity and had been strongly affected by the presence of the bacteria (Fig. 3).

**Figure 3:**
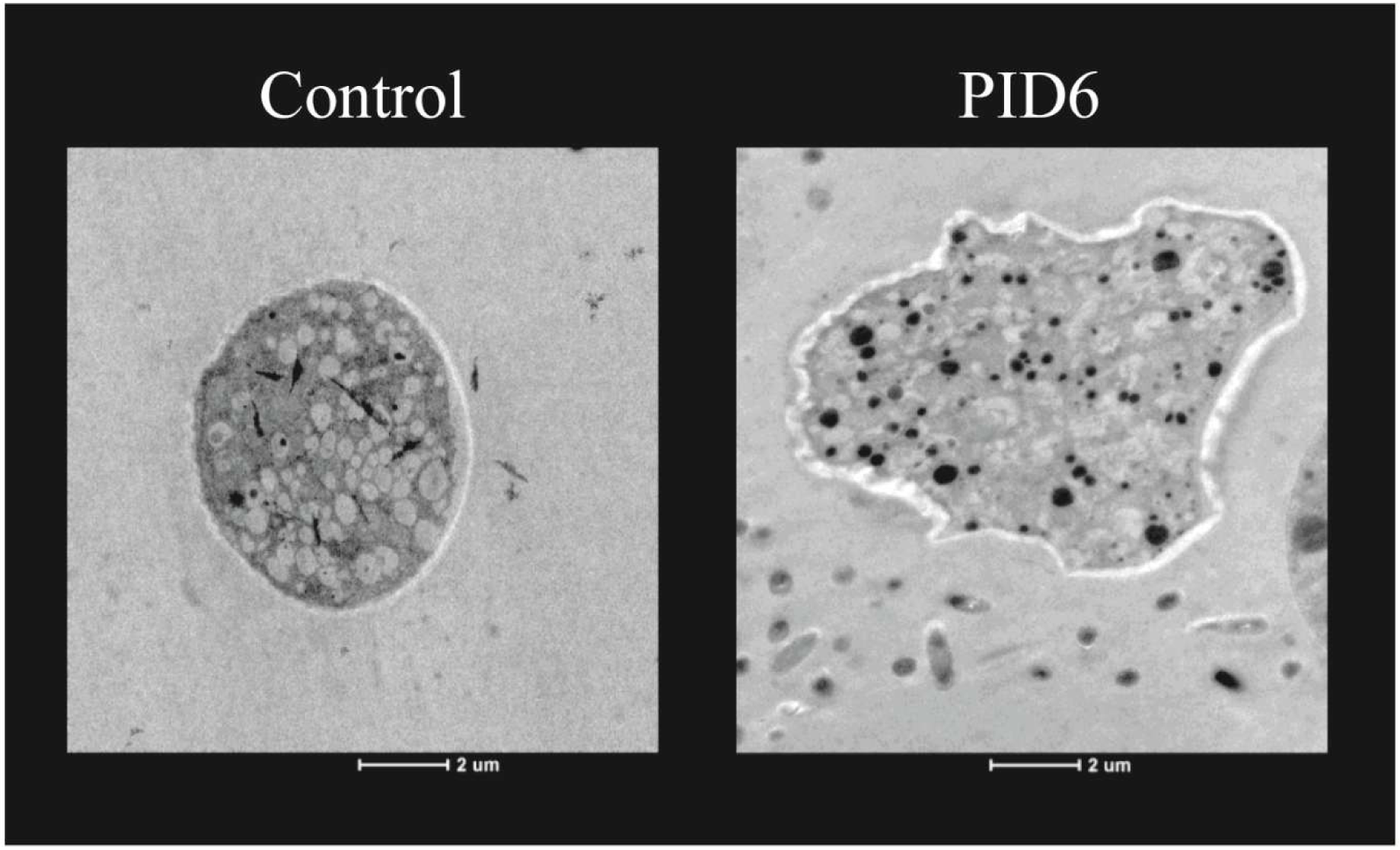
Transmission electron microscopy image of *P. viticola* zoospores placed for 30 min in contact with water (control), or a solution of the *Bacillus cereus* strain PID6 OD_600_ = 1.

### 2. Bacteria from the grapevine phyllosphere inhibit symptom development caused by *B. cinerea* and *P. viticola* infection in preventive leaf discs assays

In this next experimental part, we tested if the inoculation of bacteria in plant tissue prior to infection could reduce the development of symptoms caused by *B. cinerea* and *P. viticola* applied directly on leaf tissue. The protection conferred by bacteria was tested by inoculating leaf discs before infecting them with either pathogen.

To assess their protective activity, strains were applied to leaf discs before infection with *B. cinerea* and *P. viticola* to let them adapt to their new environment. In leaf discs infected with *B. cinerea*, preventive bacterial treatment with 15 strains reduced necrosis significantly by 38 to 81 % on average (Table 2, Fig. 4). In our conditions, seven strains, with a majority of epiphytic bacteria were able to achieve a better reduction in symptoms (more than 61 %) than the Botector® (*Aureobasidium pullulans*) used as positive control: CHP29 (*Microbacterium proteolyticum*), SOP47 (*Variovorax paradoxus*), CHP2 (*Cupriavidus metallidurans*), PIP6 (*Herbaspirillum huttiense*), PIP36 (*Bacillus cereus*), SOP8 (*Cupriavidus metallidurans*) and PID2 (*Bacillus simplex*) (Fig. 4).

**Table 2:**
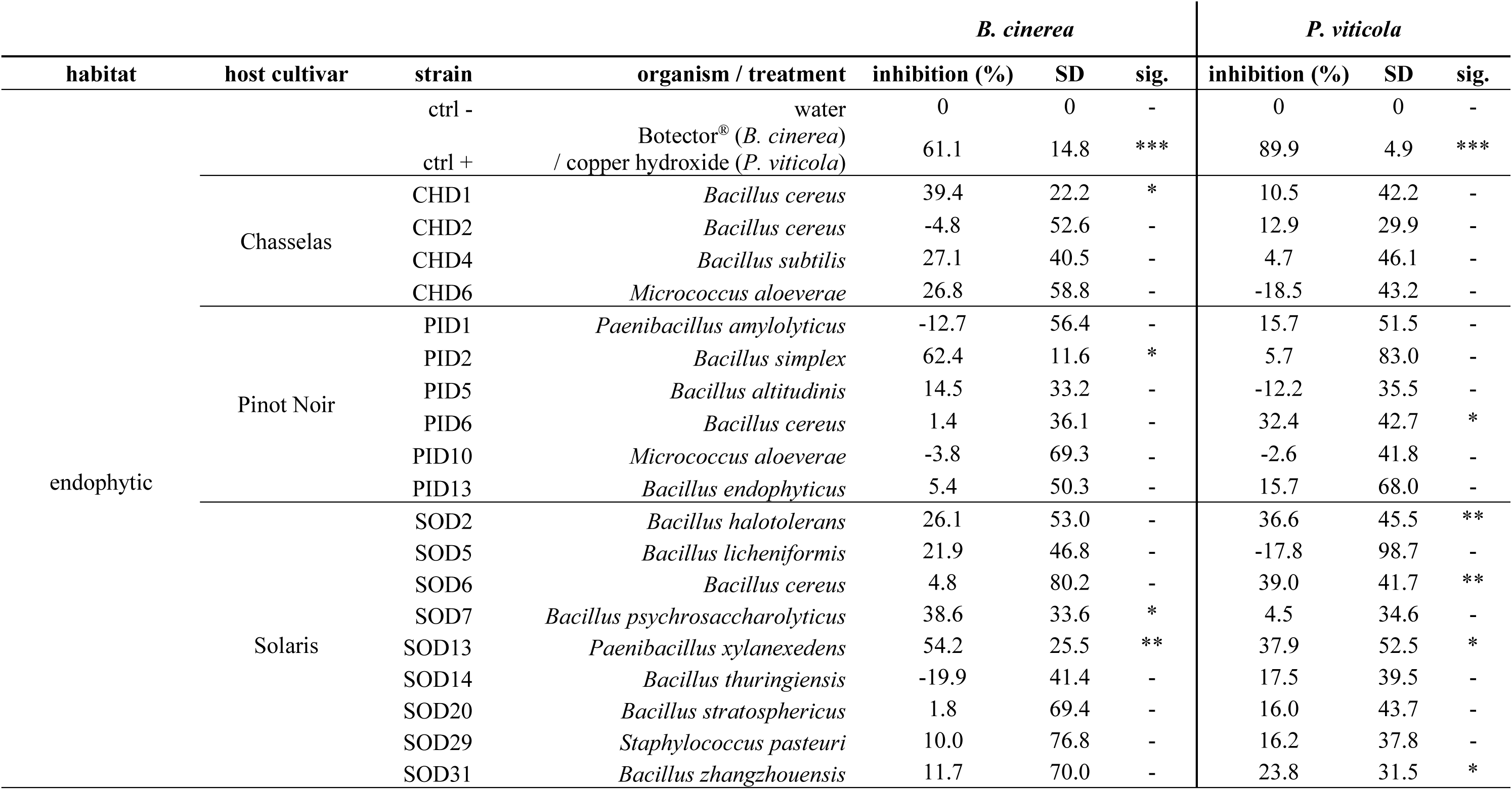
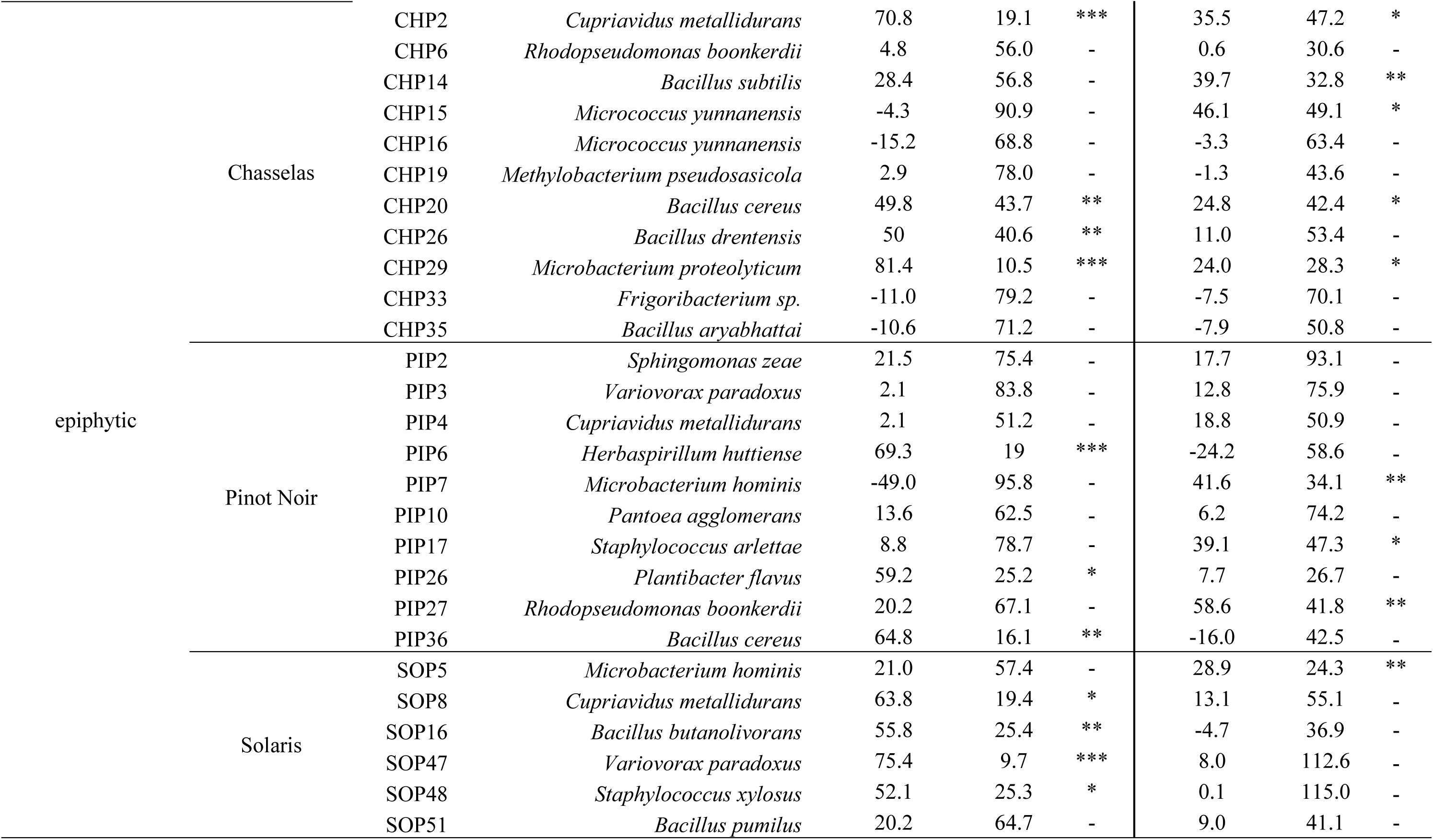
Effect of grapevine-associated phyllosphere bacteria on *B. cinerea* and *P. viticola* symptom inhibition one week after infection in grapevine leaf discs inoculated with bacteria two days before infection. Activity against *B. cinerea* is reported as the reduction of the area covered by the necrosis compared to the infected water control. Activity against *P. viticola* is reported as the reduction of the area covered by the mycelium compared to the infected water control. Statistical significance: * = *p* < 0.05; ** = *p* < 0.01; *** = *p* < 0.001; - not significant.

| habitat | host cultivar | strain | organism / treatment | *B. cinerea* |  |  | *P. viticola* |  |  |
| --- | --- | --- | --- | --- | --- | --- | --- | --- | --- |
|  |  |  |  | inhibition (%) | SD | sig. | inhibition (%) | SD | sig. |
| endophytic |  | ctrl - | water | 0 | 0 | - | 0 | 0 | - |
|  |  |  | Botector® (*B. cinerea*) |  |  |  |  |  |  |
|  |  | ctrl + | / copper hydroxide (*P. viticola*) | 61.1 | 14.8 | *** | 89.9 | 4.9 | *** |
|  | Chasselas | CHD1 | *Bacillus cereus* | 39.4 | 22.2 | * | 10.5 | 42.2 | - |
|  |  | CHD2 | *Bacillus cereus* | -4.8 | 52.6 | - | 12.9 | 29.9 | - |
|  |  | CHD4 | *Bacillus subtilis* | 27.1 | 40.5 | - | 4.7 | 46.1 | - |
|  |  | CHD6 | *Micrococcus aloeverae* | 26.8 | 58.8 | - | -18.5 | 43.2 | - |
|  | Pinot Noir | PID1 | *Paenibacillus amylolyticus* | -12.7 | 56.4 | - | 15.7 | 51.5 | - |
|  |  | PID2 | *Bacillus simplex* | 62.4 | 11.6 | * | 5.7 | 83.0 | - |
|  |  | PID5 | *Bacillus altitudinis* | 14.5 | 33.2 | - | -12.2 | 35.5 | - |
|  |  | PID6 | *Bacillus cereus* | 1.4 | 36.1 | - | 32.4 | 42.7 | * |
|  |  | PID10 | *Micrococcus aloeverae* | -3.8 | 69.3 | - | -2.6 | 41.8 | - |
|  |  | PID13 | *Bacillus endophyticus* | 5.4 | 50.3 | - | 15.7 | 68.0 | - |
|  | Solaris | SOD2 | *Bacillus halotolerans* | 26.1 | 53.0 | - | 36.6 | 45.5 | ** |
|  |  | SOD5 | *Bacillus licheniformis* | 21.9 | 46.8 | - | -17.8 | 98.7 | - |
|  |  | SOD6 | *Bacillus cereus* | 4.8 | 80.2 | - | 39.0 | 41.7 | ** |
|  |  | SOD7 | *Bacillus psychrosaccharolyticus* | 38.6 | 33.6 | * | 4.5 | 34.6 | - |
|  |  | SOD13 | *Paenibacillus xylanexedens* | 54.2 | 25.5 | ** | 37.9 | 52.5 | * |
|  |  | SOD14 | *Bacillus thuringiensis* | -19.9 | 41.4 | - | 17.5 | 39.5 | - |
|  |  | SOD20 | *Bacillus stratosphericus* | 1.8 | 69.4 | - | 16.0 | 43.7 | - |
|  |  | SOD29 | *Staphylococcus pasteurii* | 10.0 | 76.8 | - | 16.2 | 37.8 | - |
|  |  | SOD31 | *Bacillus zhangzhouensis* | 11.7 | 70.0 | - | 23.8 | 31.5 | * |
| epiphytic | Chasselas | CHP2 | *Cupriavidus metallidurans* | 70.8 | 19.1 | *** | 35.5 | 47.2 | * |
|  |  | CHP6 | *Rhodopseudomonas boonkerdii* | 4.8 | 56.0 | - | 0.6 | 30.6 | - |
|  |  | CHP14 | *Bacillus subtilis* | 28.4 | 56.8 | - | 39.7 | 32.8 | ** |
|  |  | CHP15 | *Micrococcus yunnanensis* | -4.3 | 90.9 | - | 46.1 | 49.1 | * |
|  |  | CHP16 | *Micrococcus yunnanensis* | -15.2 | 68.8 | - | -3.3 | 63.4 | - |
|  |  | CHP19 | *Methylobacterium pseudosasicola* | 2.9 | 78.0 | - | -1.3 | 43.6 | - |
|  |  | CHP20 | *Bacillus cereus* | 49.8 | 43.7 | ** | 24.8 | 42.4 | * |
|  |  | CHP26 | *Bacillus drentensis* | 50 | 40.6 | ** | 11.0 | 53.4 | - |
|  |  | CHP29 | *Microbacterium proteolyticum* | 81.4 | 10.5 | *** | 24.0 | 28.3 | * |
|  |  | CHP33 | *Frigoribacterium sp.* | -11.0 | 79.2 | - | -7.5 | 70.1 | - |
|  |  | CHP35 | *Bacillus aryabhattai* | -10.6 | 71.2 | - | -7.9 | 50.8 | - |
|  | Pinot Noir | PIP2 | *Sphingomonas zeae* | 21.5 | 75.4 | - | 17.7 | 93.1 | - |
|  |  | PIP3 | *Variovorax paradoxus* | 2.1 | 83.8 | - | 12.8 | 75.9 | - |
|  |  | PIP4 | *Cupriavidus metallidurans* | 2.1 | 51.2 | - | 18.8 | 50.9 | - |
|  |  | PIP6 | *Herbaspirillum huttiense* | 69.3 | 19 | *** | -24.2 | 58.6 | - |
|  |  | PIP7 | *Microbacterium hominis* | -49.0 | 95.8 | - | 41.6 | 34.1 | ** |
|  |  | PIP10 | *Pantoea agglomerans* | 13.6 | 62.5 | - | 6.2 | 74.2 | - |
|  |  | PIP17 | *Staphylococcus arlettae* | 8.8 | 78.7 | - | 39.1 | 47.3 | * |
|  |  | PIP26 | *Plantibacter flavus* | 59.2 | 25.2 | * | 7.7 | 26.7 | - |
|  |  | PIP27 | *Rhodopseudomonas boonkerdii* | 20.2 | 67.1 | - | 58.6 | 41.8 | ** |
|  |  | PIP36 | *Bacillus cereus* | 64.8 | 16.1 | ** | -16.0 | 42.5 | - |
|  | Solaris | SOP5 | *Microbacterium hominis* | 21.0 | 57.4 | - | 28.9 | 24.3 | ** |
|  |  | SOP8 | *Cupriavidus metallidurans* | 63.8 | 19.4 | * | 13.1 | 55.1 | - |
|  |  | SOP16 | *Bacillus butanolivorans* | 55.8 | 25.4 | ** | -4.7 | 36.9 | - |
|  |  | SOP47 | *Variovorax paradoxus* | 75.4 | 9.7 | *** | 8.0 | 112.6 | - |
|  |  | SOP48 | *Staphylococcus xylosus* | 52.1 | 25.3 | * | 0.1 | 115.0 | - |
|  |  | SOP51 | *Bacillus pumilus* | 20.2 | 64.7 | - | 9.0 | 41.1 | - |

**Figure 4:**
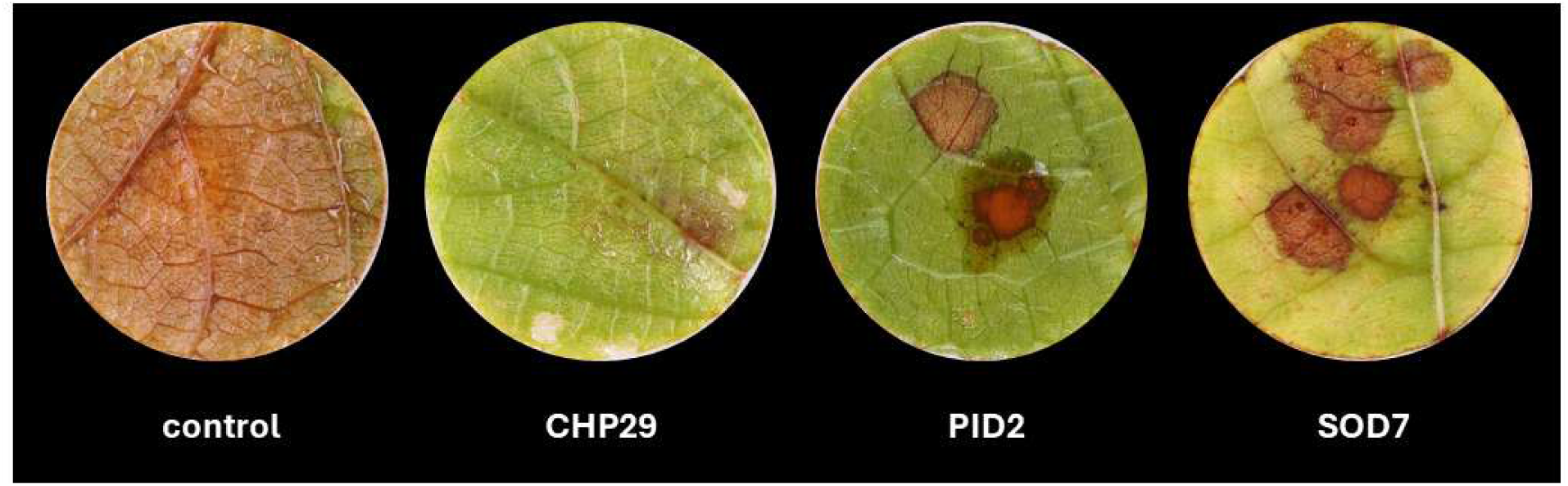
Representative pictures of grapevine leaf discs inoculated with phyllosphere bacteria prior to infection with *B. cinerea*. Control: leaf disc not inoculated with bacteria but infected with *B. cinerea* spores; CHP29, PID2 and SOD7: leaf discs inoculated with *Microbacterium proteolyticum* (CHP29), *Bacillus simplex* (PID2) and *Bacillus psychrosaccharolyticus* (SOD7), respectively, prior to infection. Pictures were taken one week after infection.

In leaf discs infected with *P. viticola,* 14 strains were able to significantly reduce the disease symptoms, to a level ranging from 24 to 59 % on average (Table 2, Fig. 5), while all other strains did not result in any significant symptom reduction. These 14 strains included 7 strains already identified as active in the first infection assay carried out with exposed zoospores described above (Table 1), while seven other strains had not shown activity in the previous experiment, but revealed their anti-oomycete abilities when pre-inoculated on the leaf discs (Table 2) : CHP15 (*Micrococcus yunnanensis*), SOD6 (*Bacillus cereus)*, SOD2 (*Bacillus halotolerans*), CHP2 (*Cupriavidus metallidurans*), CHP20 (*Bacillus cereus*), CHP29 (*Microbacterium proteolyticum*) and SOD31 (*Bacillus zhangzhouensis*).

**Figure 5:**
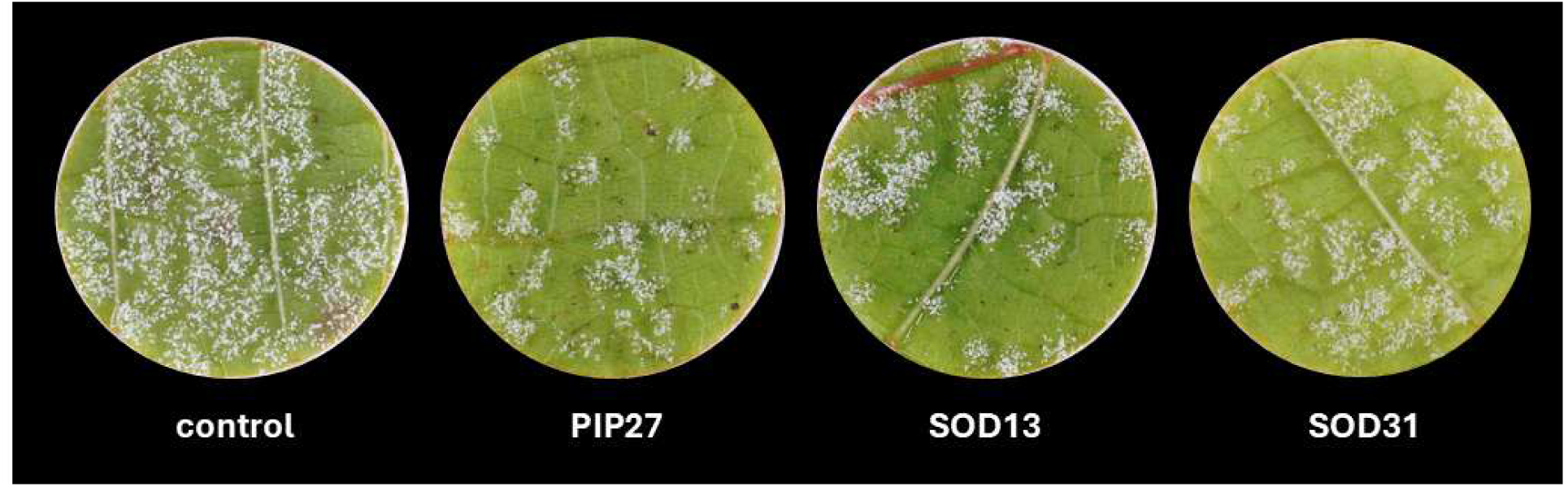
Representative pictures of grapevine leaf discs inoculated with phyllosphere bacteria prior to infection with *P. viticola*. Control: leaf disc not inoculated with bacteria but infected with *P. viticola* zoospores; PIP27, SOD13 and SOD31: leaf discs inoculated with *Rhodopseudomonas boonkerdii* (PIP27), *Paenibacillus xylanexedens* (SOD13) and *Bacillus zhangzhouensis* (SOD31), respectively, prior to infection. Pictures were taken one week after infection.

### 3. Phyllosphere bacteria stimulate the expression of several plant defense genes and promote the synthesis of several stilbenic compounds after infection

Beyond the direct effects against pathogens observed above, we next tested whether the most protective strains would also stimulate plant defenses. Infected leaf discs preinoculated with bacteria were collected to evaluate their effect on plant defenses. We monitored defense genes expression and stilbenic compound production. A set of defense genes linked to the systemic acquired resistance (PAL), the induced systemic resistance (LOX), the stilbene biosynthesis pathway (STS and ROMT), as well as two pathogenesis related protein encoding genes (PR-1 and PR-4) were used for the qPCR analysis.

For *B. cinerea*, we tested the strains that triggered an inhibition higher than that observed in the positive control. These strains exhibited diverse effects on the expression of defense-related genes, with the majority either downregulating or not significantly affecting their expression (Fig. 6). Except for the strain PIP36 (*Bacillus cereus*), which was able to promote the expression of almost all genes, SOP8 (*Cupriavidus metallidurans*), which only promoted the expression of PR-4, and ROMT, CHP29 and CHP2 which promoted the expression of ROMT and PR-1 respectively but had moderate to no effect on the other genes, all the other strains had no significant effect on the expression of the tested set of defense-related genes, suggesting that they affected the pathogen through another mechanism.

**Figure 6:**
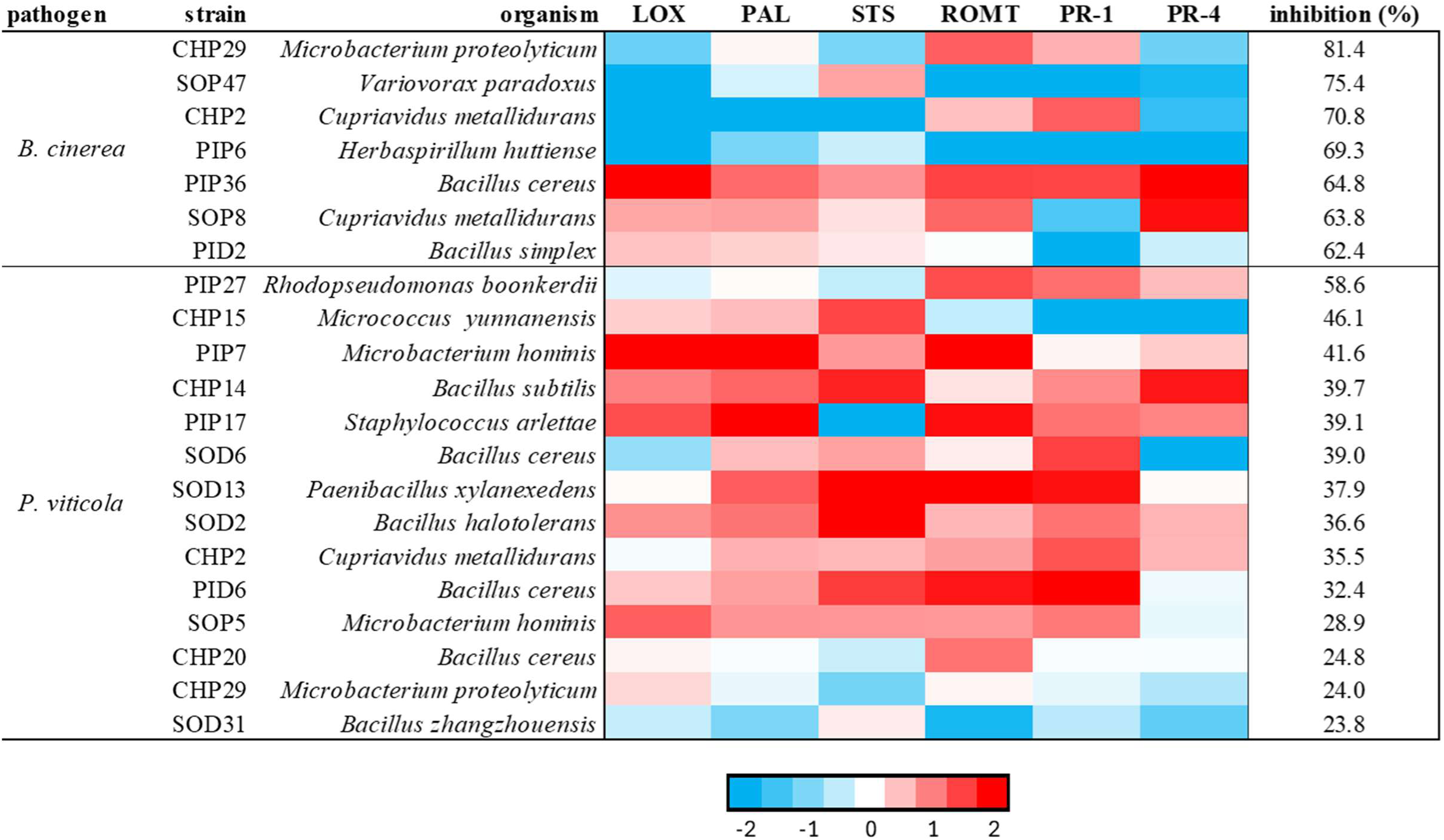
Level of expression of defense-related genes in grapevine leaf discs inoculated with phyllosphere bacteria that significantly protected the plant against *P. viticola* or *B. cinerea* in leaf disc assays. The levels of expression were normalized with the two reference genes Actin and 60SRP, and expressed relative to infected control leaf discs that were not inoculated with bacteria. Strains are ordered by their inhibition percentage, which corresponds to the reduction in symptom development caused by each strain in the leaf disc experiments.

Among the 14 strains that were effective against *P. viticola*, 11 significantly upregulated the expression of at least three defense genes and only three strains had no positive impact on the defense gene expression (Fig. 6). These latter strains (CHP20, CHP29 and SOD31), showed the lowest level of inhibition of symptoms on leaf discs and mainly downregulated gene expression or had no significant effect on the monitored genes. Among the strains that significantly upregulated the expression of several defense-genes, we observed a tendency for upregulation of the genes from the SAR and the stilbene synthesis pathways (PR-1, PAL, STS and ROMT). In contrast, the expression of the genes LOX and PR4 were induced by fewer strains. Furthermore, it seems that when these two genes were upregulated, the fold change was globally lower compared to the genes from the SAR or the stilbene pathway.

Interestingly, the strains PIP7, PIP17, SOD13, SOD2 and CHP2 were able to stimulate all or almost all genes tested without downregulating any single gene, except to PIP17 which dowregulated only STS. Finally, regarding the two best performing strains in terms of symptom reduction, PIP27 and CHP15, they had an opposite effect on gene expression: PIP27 upregulated ROMT, PR-1 and PR-4 and slightly downregulated LOX, PAL and STS, while CHP15 caused the opposite effect.

Regarding the production of stilbenic compounds, differences were only observed in plants infected with *P. viticola* (Fig. 7). In infected plants inoculated with protective bacteria, the production of piceid (an inactive storage form), resveratrol (a precursor of more active forms) and δ viniferin (an oxidized form with antimicrobial activity) was stimulated by most of the bacteria. On the contrary, the production of the active compounds, isohopeaphenol, α-viniferin and ε-viniferin was negatively impacted by the inoculation with most strains. However, few strains were able to stimulate the production of the different active forms. Accordingly, CHP15 (*Micrococcus yunnanensis*), SOD13 (*Paenibacillus xylanexedens*), SOD2 (*Bacillus halotolerans*), SOD6 and PID6 (two *Bacillus cereus*) were able to promote the synthesis of at least three of the active forms in infected leaves. Except for CHP15, which triggered an inhibition of symptoms of 46 %, these strains were responsible for an inhibition of 37, 36, 39 and 32 %, respectively, corresponding to a moderate protection level among our active strains.

**Figure 7:**
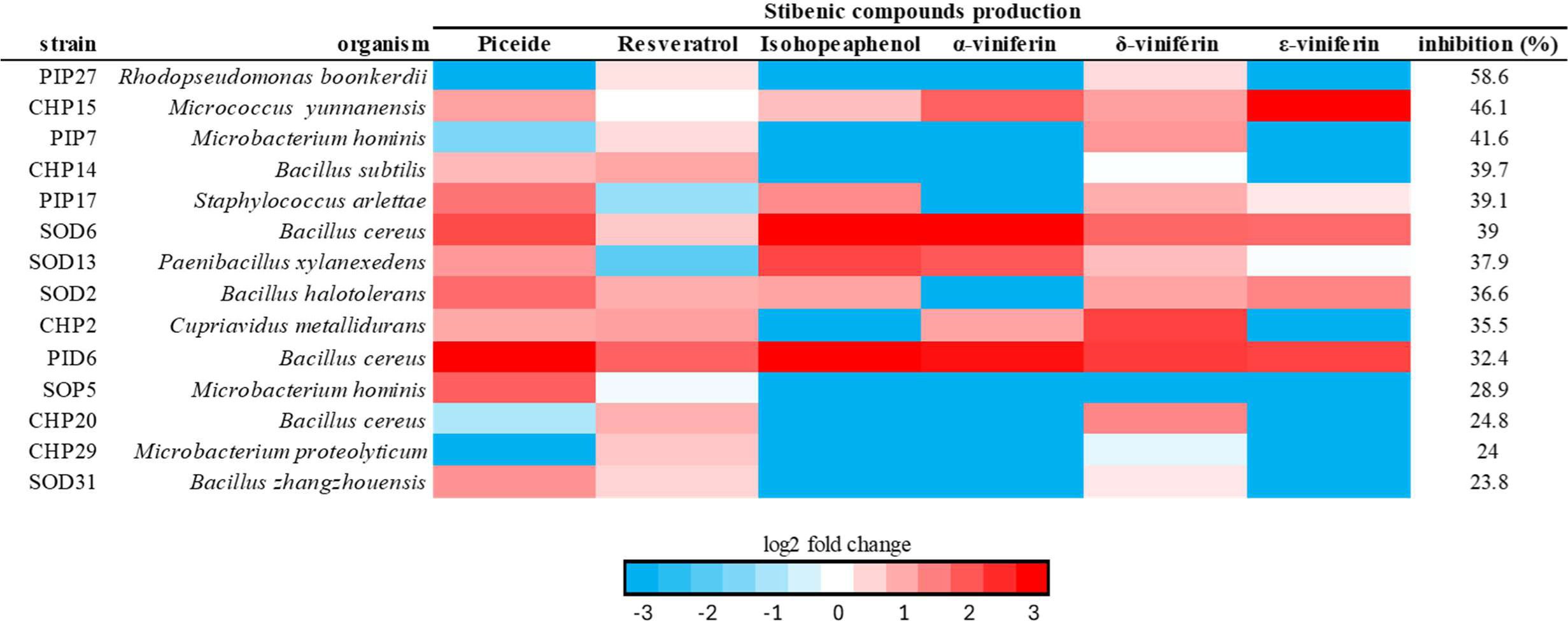
Stilbene production levels in grapevine leaf discs inoculated with phyllosphere bacteria 3 days after infection with *P. viticola* zoospores. Levels of production are based on dry weight and are expressed relative to infected leaves that were not inoculated with bacteria. Strains are ordered by their inhibition percentage, which corresponds to the reduction in symptom development achieved with each strain in leaf disc assays.

### 4. Selection of bacteria for the consortia

A selection of strains that had shown significant effects against *B. cinerea* and *P. viticola* was performed for a use in combination, based both on effectiveness and on the resistance to phytosanitary products. The goal was to build consortia of three strains which would be effective against *B. cinerea, P. viticola* and display plant defense stimulation abilities.

#### 4.1 Effectiveness threshold against *B. cinerea* and *P. viticola*

Regarding the effects on propagules *in vitro*, we selected strains able to at the same time fully inhibit the germination of at least 50 % of *B. cinerea* spores, inhibit *P. viticola* zoospore motility and provide a significant level of symptom inhibition with the subsequent infection. Eight strains met these criteria, five *Bacillus,* two *Microbacterium* spp. and one *Sphingomonas* sp. (Table 3).

**Table 3:** Summary of the selection criteria used to select bacteria consortia. Strains were evaluated for i) *in vitro* inhibition of *B. cinerea* conidial germination and *P. viticola* zoospore motility, ii) inhibition of symptoms caused by *B. cinerea* and *P. viticola* on grapevine leaf discs, and iii) resistance to biofungicide products. A cross indicates that the strain met the following selection thresholds. Propagules: at least 50% inhibition of *B. cinerea* conidial germination and inhibition of *P. viticola* zoospore motility with significant symptom reduction in the subsequent infection assay; leaf tissue: reduction of symptoms of at least 61 % for *B. cinerea* and reduction of symptoms of at least 30 % for *P. viticola*; biofungicide resistance: survival in the presence of potassium bicarbonate (0.2 %) and wettable sulfur (0.4 %), tested individually and in combination. The seven strains fulfilling the selection criteria are marked in bold.

| strain | organism / treatment | *P. viticola*<br>propagule inhibition | *B. cinerea*<br>propagule inhibition | *B. cinerea* inhibition<br>on leaf discs | *P. viticola* inhibition<br>on leaf discs | resistance to<br>biofungicides |
| --- | --- | --- | --- | --- | --- | --- |
| PID2 | *Bacillus simplex* | x | x | x |  |  |
| PID6 | ***Bacillus cereus*** | x | x |  | x | x |
| PID13 | ***Bacillus endophyticus*** | x | x |  |  | x |
| SOD2 | ***Bacillus halotolerans*** |  | x |  | x | x |
| SOD6 | *Bacillus cereus* |  | x |  | x |  |
| SOD13 | *Paenibacillus xylanexedens* |  | x |  | x |  |
| CHP2 | ***Cupriavidus metallidurans*** |  | x | x | x | x |
| CHP14 | ***Bacillus subtilis*** | x | x |  | x | x |
| CHP15 | *Micrococcus yunnanensis* |  |  |  | x |  |
| CHP29 | *Microbacterium proteolyticum* |  | x | x |  |  |
| PIP2 | *Sphingomonas zeae* | x | x |  |  |  |
| PIP6 | ***Herbaspirillum huttiense*** |  | x | x |  | x |
| PIP7 | *Microbacterium hominis* | x | x |  | x |  |
| PIP10 | *Pantoea agglomerans* | x |  |  |  |  |
| PIP17 | *Staphylococcus arlettae* | x |  |  | x |  |
| PIP27 | *Rhodopseudomonas boonkerdii* |  |  |  | x |  |
| PIP36 | *Bacillus cereus* | x | x | x |  |  |
| SOP5 | *Microbacterium hominis* | x | x |  |  |  |
| SOP8 | ***Cupriavidus metallidurans*** |  | x | x |  | x |
| SOP47 | *Variovorax paradoxus* |  | x | x |  |  |
| SOP51 | *Bacillus pumilus* | x |  |  |  |  |

For effects against *B. cinerea* symptoms on leaf discs, we only considered strains with an effectiveness higher than the 61 % reached by the positive control Botector® (*Aureobasidium pullulans*). For *P. viticola*, since the reduction of symptoms was lower in the preventive leaf disc assays, the threshold of effectiveness was set to 30 %. These criteria were met by seven strains for *B. cinerea* and by 10 strains for *P. viticola* (Table 3).

#### 4.2 Resistance to plant protection products

Even though in this study, the effects of the bacteria were tested under laboratory conditions, it is important to consider that ultimately, the bacteria would be used in the field under real agricultural conditions. They could be in contact with chemical plant protection solutions, and it is therefore important that the selected strains are not susceptible to such products. We therefore decided to select strains based on their ability to survive in presence of two biofungicides used in organic viticulture, potassium bicarbonate and wettable sulfur, alone or in combination (Table S2).

Seven strains were capable of both affecting pathogen propagules and/or providing a significant plant protection as well as of surviving in presence of the tested phytosanitary products: PID6 (*Bacillus cereus*), PID13 (*Bacillus endophyticus*), SOD2 (*Bacillus halotolerans*), CHP2 (*Cupriavidus metallidurans*), CHP14 (*Bacillus subtilis*), PIP6 (*Herbaspirillum huttiense*) and SOP8 (*Cupriavidus metallidurans*) (Table 3). No incompatibility was observed between these strains when grown together in liquid coculture, as tested by their subsequent cultivation on selective media plates. It appeared that they were all able to survive adequately when grown together in the same mixture.

### 5. Protective effects of bacteria consortia against pathogens

Combinations of the seven selected bacteria were tested against *B. cinerea* spore germination *in vitro* and *P. viticola* infection on whole plants to compare their protective effect with that of single strains.

#### 5.1. The inhibition of *B. cinerea* spore germination was higher with consortia than with bacteria alone

Consortia containing two to three of the seven selected strains were tested. The solution containing the combination of bacteria had the same final cell concentration as the one used in the previous experiment with single strains, so any increase in effectiveness could not be simply due to the increased bacterial density in the mixture. This combination resulted in an overall increase of the strains’ germination inhibition abilities. All consortia were able to repress the germination process, with the least effective one (CHP14, PIP6 and SOD2) affecting 98 % of the spores and 20 out of the 23 tested consortia affecting 100 % of the spores (Fig. 8). Consortia differed mainly in the proportion of spores that were fully vs. partially inhibited in their germination. The abilities of consortia to fully inhibit the germination of *B. cinerea* conidia were compared to the best-performing strain within each combination using the Highest Single Agent (HSA) method (Table S3). Among the 23 combinations tested, 7 displayed a reduced effectiveness (≤ 10 percentage points lower than HSA), 9 showed a similar level of inhibition (within ± 10 percentage points compared to HSA), and 7 performed better (≥ 10 percentage points higher than HSA). The highest improvements were observed in consortia of two strains containing bacteria with low to medium effects regarding full inhibition of spore germination: PID6/SOP8 and CHP2/PID6 with an increase of 67 and 82 % compared to their HSA respectively. Accordingly, three-strains consortia including these four strains like, performed similarly to, or better than their HSA. In contrast, all three strains consortia including SOD2 performed not better, and even significantly worse than their HSA.

**Figure 8:**
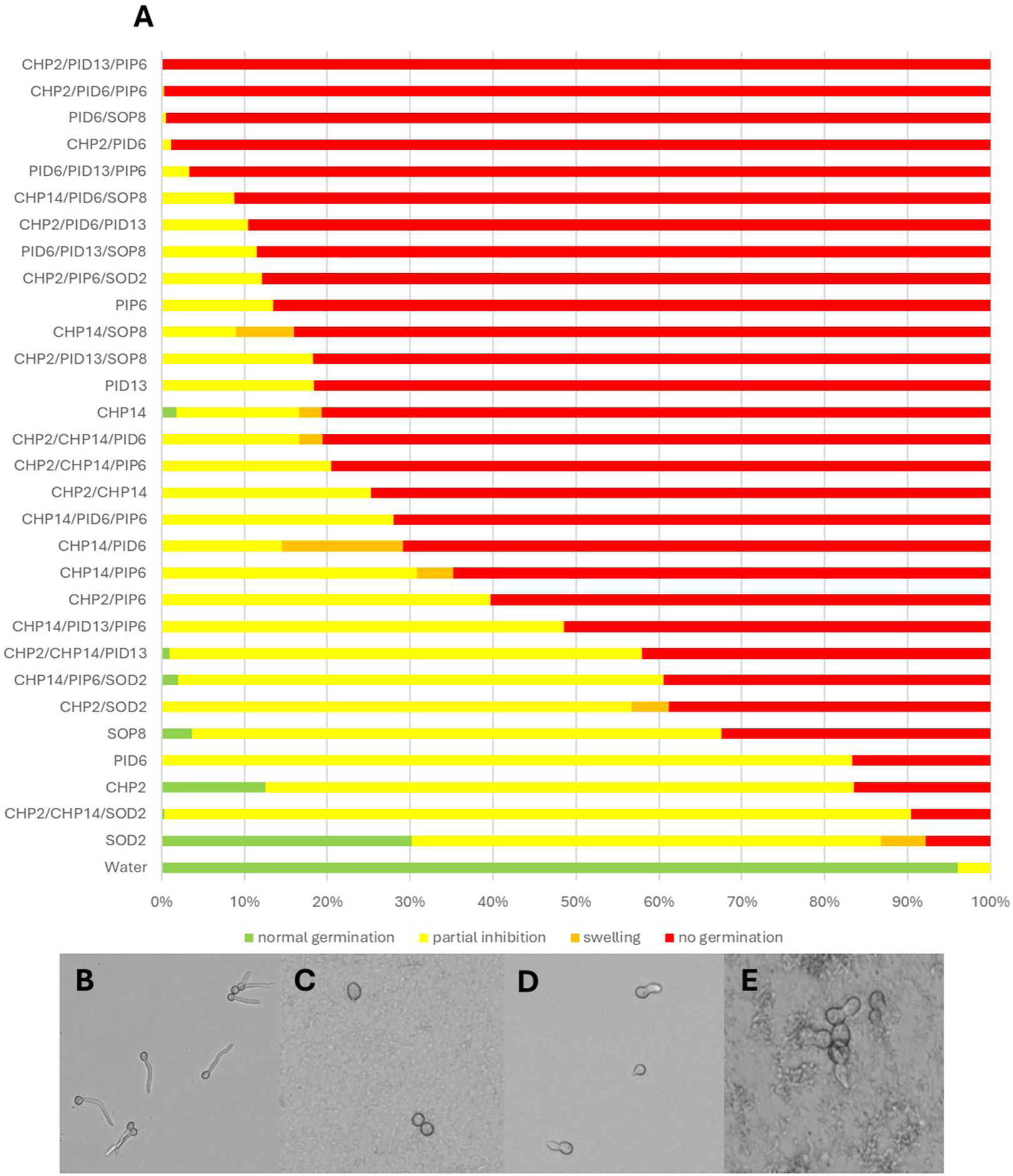
Effect of consortia of phyllosphere bacteria on *B. cinerea* spore germination. A: Each colored bar represents the proportion of spores showing the corresponding morphology. Green: normal germination; yellow; partial inhibition (only the tip of the germ tube is visible); orange: swelling and red: no germination. Each treatment with bacteria significantly affected the physiology of the spores, with *p* < 0.05 according to a Fisher’s exact test with Bonferroni’s correction. The different types of germination morphologies are shown in (B to E); B: normal germination; C: absence of germination; D: partial germination and E: swelling.

#### 5.2 Strains combinations were more efficient and consistent than single strains for whole plant protection against *P. viticola*

The seven selected strains were assembled to build four consortia each containing at least one strain capable of impairing *B. cinerea* and *P. viticola* propagules, inhibiting symptom development of both pathogens, and stimulating plant defenses. Consortia were sprayed on half the leaves of grapevine plants (leaf stages 4 to 7), while the other half was sprayed with water prior to whole plant infection with *P. viticola* zoospores. After a week, symptoms were measured on bacterial-inoculated and uninoculated leaves in search of local and systemic protective effects of the bacteria. Results showed that the strains CHP14, CHP2 and PID13 alone provided the lowest reduction of symptoms along with the highest variability (38, 34 and 31 % respectively), the reduction triggered by CHP2 and PID13 being not even significant. However, all combinations were able to significantly reduce the level of symptoms except for SOP8/CHP14/PID6 (Fig. 9 and Table 4). For all plants and for all treatments except for PIP6, the severity of symptoms was statistically similar on bacteria-inoculated leaves and in leaves that were sprayed with only water, suggesting that the presence of bacteria triggered a systemic response. However, the level of symptom reduction was different between the different treatments. The three best combinations (CHP2/PID6/PIP6; CHP14/PID13/PIP6 and CHP2/CHP14/PID13) led to a symptom reduction ranging from 88 to 94 % on average, reaching an almost complete protection, however still lower than the positive control (Table 4). The combination (CHP2/CHP14/SOD2) provided a reduction of 58 % which, although significant, was not very consistent (standard deviation of 39.2) and remained lower than the protection generated by the three best single strains PID6, SOP8 and PIP6 alone (82, 68 and 63 % respectively). The last combination (SOP8/CHP14/PID6) did not cause a significant reduction of symptoms either locally and systemically despite the fact that the single strains PID6 and SOP8 are the most effective strains.

**Figure 9:**
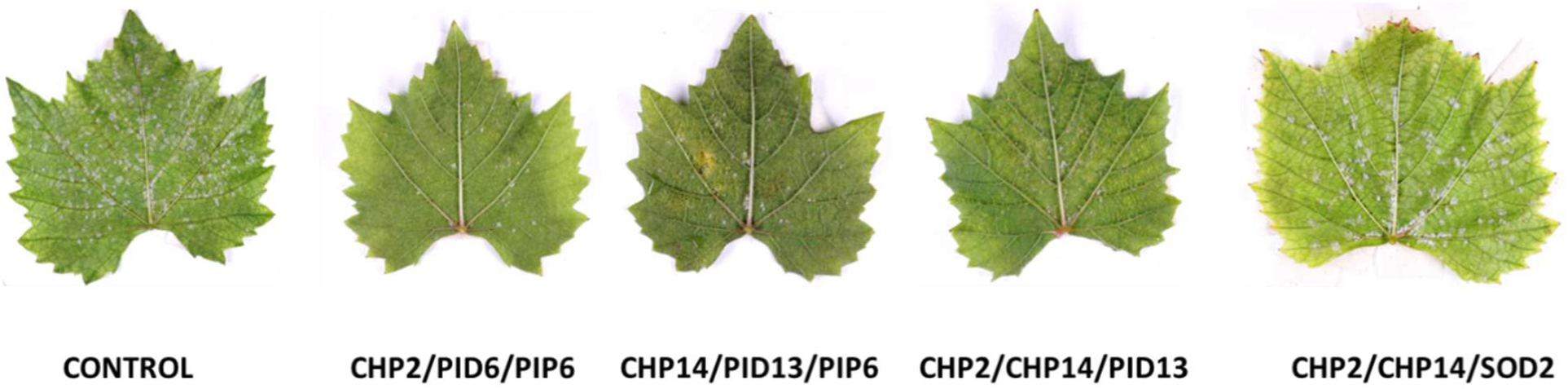
Representative pictures of grapevine leaves from whole grapevine plants inoculated with phyllosphere bacteria prior to infection with *P. viticola*. Leaves were inoculated with different mixtures of the following six strains: *Cupriavidus metallidurans* (CHP2), *Bacillus cereus* (PID6), *Herbaspirillum huttiense* (PIP6), *Bacillus subtilis* (CHP14), *Bacillus endophyticus* (PID13) and *Bacillus halotolerans* (SOD2). Control: non-inoculated leaf infected with *P. viticola* zoospores. Pictures were taken one week after infection.

**Table 4:**
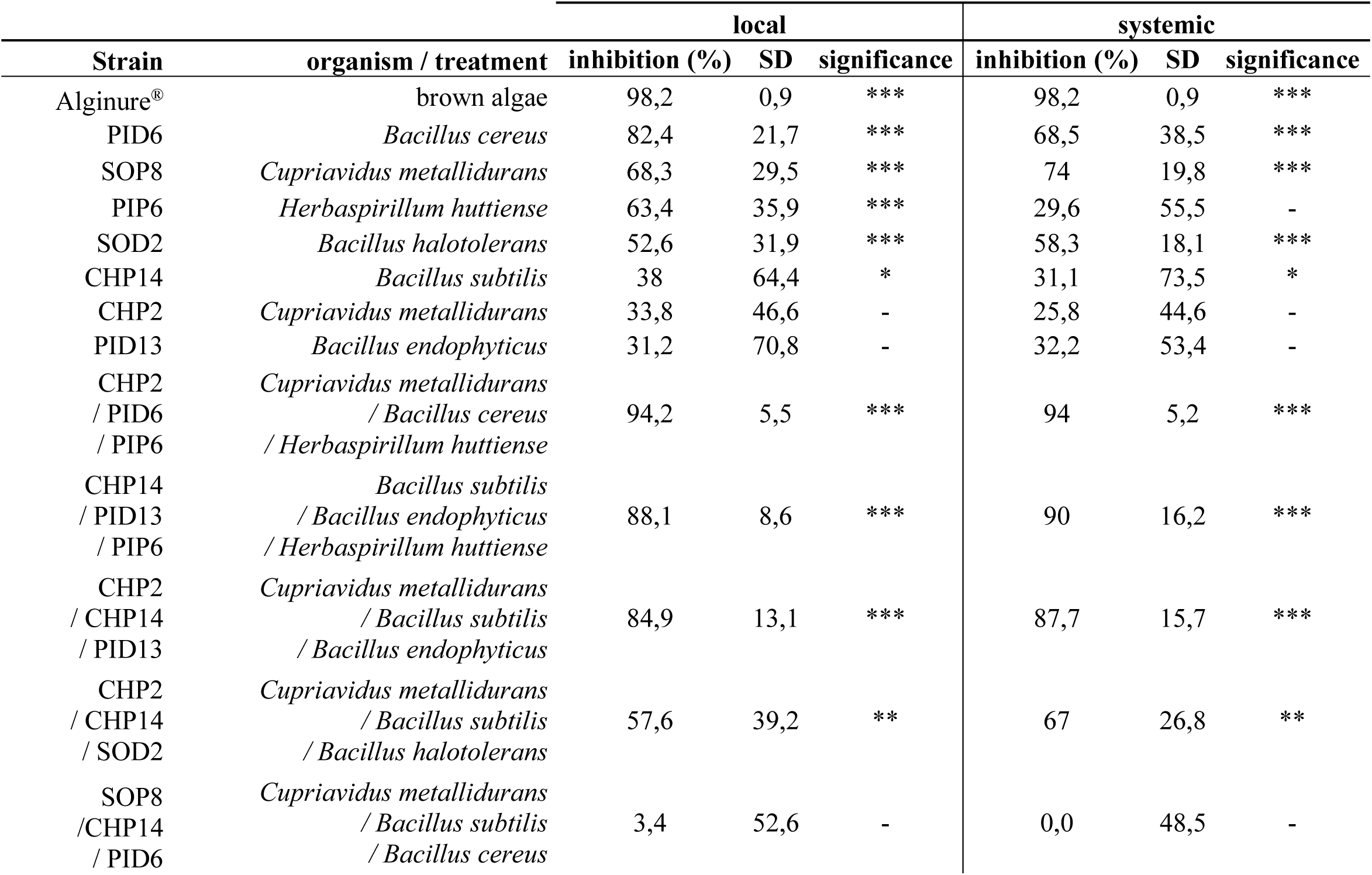
Local and systemic *P. viticola* symptom inhibition one week after infection in whole plants pre-inoculated with phyllosphere bacteria relative to uninoculated control. Statistical significance: * = p < 0.05; ** = p < 0.01; *** = p < 0.001; - not significant.

| Strain | organism / treatment | local |  |  | systemic |  |  |
| --- | --- | --- | --- | --- | --- | --- | --- |
|  |  | inhibition (%) | SD | significance | inhibition (%) | SD | significance |
| Alginure® | brown algae | 98,2 | 0,9 | *** | 98,2 | 0,9 | *** |
| PID6 | *Bacillus cereus* | 82,4 | 21,7 | *** | 68,5 | 38,5 | *** |
| SOP8 | *Cupriavidus metallidurans* | 68,3 | 29,5 | *** | 74 | 19,8 | *** |
| PIP6 | *Herbaspirillum huttiense* | 63,4 | 35,9 | *** | 29,6 | 55,5 | - |
| SOD2 | *Bacillus halotolerans* | 52,6 | 31,9 | *** | 58,3 | 18,1 | *** |
| CHP14 | *Bacillus subtilis* | 38 | 64,4 | * | 31,1 | 73,5 | * |
| CHP2 | *Cupriavidus metallidurans* | 33,8 | 46,6 | - | 25,8 | 44,6 | - |
| PID13 | *Bacillus endophyticus* | 31,2 | 70,8 | - | 32,2 | 53,4 | - |
| CHP2 | *Cupriavidus metallidurans* |  |  |  |  |  |  |
| / PID6 | / *Bacillus cereus* | 94,2 | 5,5 | *** | 94 | 5,2 | *** |
| / PIP6 | / *Herbaspirillum huttiense* |  |  |  |  |  |  |
| CHP14 | *Bacillus subtilis* |  |  |  |  |  |  |
| / PID13 | / *Bacillus endophyticus* | 88,1 | 8,6 | *** | 90 | 16,2 | *** |
| / PIP6 | / *Herbaspirillum huttiense* |  |  |  |  |  |  |
| CHP2 | *Cupriavidus metallidurans* |  |  |  |  |  |  |
| / CHP14 | / *Bacillus subtilis* | 84,9 | 13,1 | *** | 87,7 | 15,7 | *** |
| / PID13 | / *Bacillus endophyticus* |  |  |  |  |  |  |
| CHP2 | *Cupriavidus metallidurans* |  |  |  |  |  |  |
| / CHP14 | / *Bacillus subtilis* | 57,6 | 39,2 | ** | 67 | 26,8 | ** |
| / SOD2 | / *Bacillus halotolerans* |  |  |  |  |  |  |
| SOP8 | *Cupriavidus metallidurans* |  |  |  |  |  |  |
| /CHP14 | / *Bacillus subtilis* | 3,4 | 52,6 | - | 0,0 | 48,5 | - |
| / PID6 | / *Bacillus cereus* |  |  |  |  |  |  |

## Discussion

The overall results of this study indicate that a significant number of phyllosphere bacteria have complementary multi-level properties that significantly reduced the symptoms of two diseases affecting grapevine. When assembled into consortia, the strains provided stronger and more consistent protection of the plant.

### Disruption of spore germination in *B. cinerea* and of zoospore motility in *P. viticola*

The first layer of protection consists in affecting the spores, the pathogens’ unit of dispersion and infection, which will prevent the infectious process. In our study, among the 28 strains able to impair the motility of *P. viticola* zoospores, almost 20 strains belonged to the genus *Bacillus*. This genus is well known for containing many biocontrol species and several studies have highlighted the effect of *Bacillus* strains on zoospores. *Bacillus* sp. 109GG020 and *B. licheniformis* were shown to affect *Phytophthora capsici* zoospore motility (Tareq et al., 2015; Y. Li et al., 2020). Comparable effects have been described using *Bacillus amyloliquefaciens* against *Phytophthora sojae* (D. Liu et al., 2019) and *Bacillus cereus* against the pathogen *Pythium torulosum* (Shang et al., 1999). However, in our study, except only two *Bacillus* strains (PID6 and PID2), the most active strains which, after co-incubation with the zoospores, induced a reduction in the subsequent development of symptoms, belonged to the *Pantoea*, *Microbacterium* and *Rhodopseudomonas* genera which have so far not been reported to harbor strains affecting zoospores of oomycetes (Mondol et al., 2017). In our work, no abnormal swimming was observed after contact with the bacteria, and we did not observe any decrease in the number of retrieved zoospores compared with the non-treated controls. However, the TEM observations of zoospores exposed to the most effective strain (*B. cereus* PID6) showed that the zoospores were strongly affected since they displayed a complete loss of their typical oval shape, suggesting a severe loss of integrity. Despite the strong effects of several bacteria on zoospore motility, the surviving zoospores were still able to infect grapevine leaf discs, although with a significant reduction in symptoms compared to the control. Motility is an essential function of the zoospore for plant infection in oomycetes (Chepsergon et al., 2020). We therefore expected that zoospores unable to swim would be less likely to infect plants. However, several zoospores were still able to cause infection despite this loss of function. It seems that they were still able to reach an infection site. Thus, zoospores treated with strains like SOD2 and CHP14, despite an almost full loss of motility, were still able to successfully infect the plants. In contrast, some zoospores treated with strains such as PIP27 and PIP4 that did not inhibit spore motility showed a limited capacity to infect plants. These results suggest that loss of motility alone is not enough to fully hinder the infection of *P. viticola* zoospores and that the contact with bacteria not only affects the motility but also other key functions in the zoospores, as highlighted by the TEM observation (Fig. 3).

The phyllosphere bacteria not only affected the spore motility of *P. viticola* but also the spore germination of *B. cinerea*. The most effective strains regarding this effect belonged to the genera *Herbaspirillum*, *Cupriavidus* and *Bacillus.* The biocontrol abilities for these two first genera have not been well studied so far, but the effect of *Bacillus* strains on fungal pathogens including *B. cinerea* is well known. *Bacillus subtilis* has been reported to affect *B. cinerea* hyphal morphology and to inhibit spore germination through the release of antifungal volatiles (H. Chen et al., 2008). The use of a broth from *B. cereus* strains also caused comparable effects (Li et al., 2012), as did the application of *B. mycoides* (from the *B. cereus* group) as a biocontrol agent against *B. cinerea* in strawberry (Guetsky et al., 2001a). Interestingly, one strain of *B. thuringiensis* (SOD14) was found among the four best performing *Bacillus* strains in the present study. This species is also a member of the *B. cereus* group and a source of powerful biocontrol agents against insects, mainly due to the production of Cry and Cyt toxins by several of its members (Bravo et al., 2011), but also because of their production of chitinase (Honda et al., 2017). Since *B. cinerea* hyphae and spores are also composed of chitin (Chardonnet et al., 1999), chitinases could be responsible for this germination inhibition. Indeed, *B. thuringiensis* strains were recently suggested to be very effective biocontrol agent against fungal pathogens (Marinez-Zavala et al., 2024). Beyond Cry and Cyt toxins, *Bacilli* produce other compounds with antifungal or anti-oomycete activity like cyclic lipopeptides harboring membrane disruption abilities, such as iturin, bacillomycin and fengycin. These compounds could also be involved in the germination disruption capacity of *B. thuringiensis* (SOD14) as well as its potency to inhibit successfully the motility of *P. viticola*(Balleux et al., 2025; Markelova & Chumak, 2025; Romero et al., 2007).

### Preventive inoculation of phyllosphere bacteria on leaf discs prior to infection led to significant protection against *P. viticola* and *B. cinerea*

When applied on leaves two days prior infection, fewer bacterial strains caused a significant reduction of symptoms against *P. viticola* compared with when bacteria were mixed with zoospores prior to leaf inoculation. This reduction in the number of effective strains is probably due to the infection method; since zoospores were deposited directly on the leaf tissue and without preexposure to bacteria, their motility was still intact. Moreover, in this setup, zoospores may have started to infect the leaves before being impaired by bacteria. If we compare the list of effective strains from the first assay that were at the same time able to affect motility and/or to reduce symptom development, with the strains effective in the bacterial preinoculation experiment, seven strains, SOD13, PID6, SOP5, PIP7, PIP17, PIP27 and CHP14 (*Paenibacillus xylanexedens*, *Bacillus cereus*, two strains of *Microbacterium hominis, Staphylococcus arlettae, Rhodospeudomonas boonkerdii* and *Bacillus subtilis* respectively) proved effective in both experimental setups. Once again, the genera from the previous experiment *i.e.*: *Cupriavidus, Paenibacillus Rhodopseudomonas, Micrococcus* and a large representation of *Bacillus* species were found to be active, but we also found three different strains of *Microbacterium* among active strains. So far, this genus has not been often mentioned in the literature as effective against *P. viticola,* but some strains have been shown to stimulate plant defense and inhibit the growth of oomycetes and several fungal pathogens (Barnett et al., 2006; Freitas et al., 2019; Ray et al., 2021; Suárez-Estrella et al., 2023).

The same observations were made with *B. cinerea*: fewer strains caused a reduction in symptoms compared to the test performed on spores, suggesting that for both oomycetes and fungi, activities on spores, although helpful, are not sufficient for achieving effective biocontrol. All genera containing strains active against *P. viticola,* except *Micrococcus*, also exhibited activity against *B. cinerea*, with the addition of one *Herbaspirillum* (PIP6), one *Variovorax* (SOP47) and one *Plantibacter* (PIP26). Strains belonging to these two last genera are known for their plant growth promoting abilities (Giannelli et al., 2024; Han et al., 2013; Mayer et al., 2019; Sun et al., 2018), but their biocontrol abilities are poorly studied. *Variovorax* strains have shown moderate biocontrol activities in tomatoes against *B. cinerea* and in wheat against *Fusarium* (Besset-Manzoni et al., 2019; Köhl et al., 2020), but to our knowledge, no study has yet reported any biocontrol effects of *Plantibacter* strains.

### Several phyllosphere bacteria contribute to enhance grapevine defenses

Bacterial strains successfully inhibiting the development of symptoms when they were inoculated in leaf discs prior to infection were tested for their ability to stimulate the expression of several genes belonging to different defense pathways. Most of the strains effective against *P. viticola* were able to upregulate multiple defense genes, but several bacteria such as CHP20, CHP29 and SOD31 showed little to no significant effect regarding the selected defense gene upregulation and seemed to even slightly downregulate plant defenses. Most of the strains shared the downregulation of the PR-4 gene, a gene encoding a protein with chitinase and chitin-binding activities (Enoki & Suzuki, 2016; M. Y. Li et al., 2020; G. T. Liu et al., 2021). The expression of this protein has been shown to be upregulated in grapevine cultivars that are resistant to *P. viticola* or in susceptible cultivar upon transgenic overexpression, but not in wild-type susceptible cultivars (Enoki & Suzuki, 2016; M. Y. Li et al., 2020; G. T. Liu et al., 2021). For most bacteria stimulating plant defenses against *P. viticola*, the genes that were most frequently upregulated, PAL, STS, ROMT and PR1, were all markers of the SA pathway as expected against a biotrophic pathogen (J. Y. Chen et al., 2006; Chong et al., 2008). Interestingly, it seems that the most effective strains against *P. viticola*, PIP27 and CHP15, were not the best at stimulating plant defenses, so we may assume that they have a direct and strong effect on the pathogen.

When testing for induction of defense genes upon infection with *B. cinerea*, we observed that except for PIP36 and SOP8 that triggered the upregulation of LOX, PR-4 and ROMT (which is consistent with the results obtained with other strains by Trusov et al., 2009 and Van Loon et al., 2006), the other phyllosphere isolates negatively affected the expression of most of the defense-related genes we monitored. If we compare this observation with the results obtained for *P. viticola*, we observe that *Cupriavidus metallidurans* CHP2 stimulated the expression of defense-related genes in presence of *P. viticola*, but mostly inhibited the expression of defense-related genes in presence of *B. cinerea*. Yet, the reduction in plant defense did not appear to be detrimental to the plant, since despite the lack of plant defense stimulation, the bacteria were able to successfully protect the plant against the pathogen, suggesting that their effect was more likely acting on the pathogen rather than stimulating plant defenses. (Zhu et al., 2019; Rahman et al., 2024)

The inoculation of bacteria also affected the production of stilbenes in leaves. These compounds are phytoalexins with different levels of antifungal properties depending on their structure (Viret et al., 2018). In plants infected with *P. viticola*, multiple strains were able to stimulate the production of several of these compounds. Most of the strains promoted the production of resveratrol, synthesized by the stilbene synthase (STS), an active form of stilbene and precursor of several more active compounds such as viniferins and pterostilbenes (Coutos-Thévenot et al., 2001). CHP15, SOD13, CHP2, SOD6 and PID6 (*M. yunnanensis*, *B. xylanexedens*, *C. metallidurans* and two *B. cereus* strains) were able to strongly increase the production of at least two different forms of viniferins, one of the most active stilbenic compounds against *P. viticola* (Alonso-Villaverde et al., 2011; Pezet et al., 2003; Van Leeuwen et al., 2013). The correlation between the increased resveratrol and viniferin production and the higher expression of STS genes induced by several active strains suggests that the protection mediated by these strains is indeed mediated, at least partly, by the induction of this pathway. Similar effects of beneficial microbes on the grapevine stilbene biosynthesis pathway have been described in Rühmann et al. (2013) with the yeast *Aureobasidium pullulans*. In contrast to viniferins, we could not detect pterostilbenes, which are compounds with higher antifungal effects than viniferins, in any sample, despite the increase in the expression of the ROMT gene encoding the enzyme responsible for their synthesis from resveratrol.

On the contrary, no significant increase of any stilbenic compound was found upon *B. cinerea* infection, but these compounds have not yet been shown to be effective against *B. cinerea* dormant conidia (Pezet & Pont, 1990; Pont & Pezet, 1990).

### Combinations of phyllosphere bacteria are more effective than single strains in protecting grapevine plants against *P. viticola*

The combination of strains selected based on their complementary effects on pathogen spores, symptom development and plant defenses provided high protection with low variability on whole plants. This protection was observed at comparable levels both locally and systemically (Table 4). Interestingly, as observed in experiments targeting *B. cinerea* spore germination (Fig. 8), the combination of strains containing three different genera was the most effective one against *P. viticola* compared to the other combinations containing at least two *Bacillus* strains. The combination CHP2/PID6/PIP6 (*Cupriavidus metallidurans*, *Bacillus cereus* and *Herbaspirillum huttiense*) was the most effective one and was able to reach an almost full inhibition of symptoms (94 %) with very little variability (SD 5.5). The best level of protection (98 %) was obtained by the positive control Alginure®, a product certified for field use against *P. viticola* in Switzerland containing a combination of brown algae, potassium phosphonate and amino acids, acting as a direct fungicide and as a plant defense enhancer. Our best combination of bacteria, despite the absence of any formulation, was therefore able to match the protection conferred by a mixture of biocontrol and chemical agents. However, two combinations performed less well than expected. The CHP2/CHP14/SOD2 consortium conferred a weaker reduction in symptoms than the three best single-strains. This result is consistent with our observations on *B. cinerea* spores, where the presence of the strain SOD2 (*Bacillus halotolerans*) seemed to limit the performance of consortia in which it was included. Finally, the weakest consortium, SOP8/CHP14/PID6, showed no visible reduction in symptoms despite containing two of the best single strains (SOP8 and PID6). This consortium also exhibited the highest variability of all combinations.

In the whole plant assay against *P. viticola*, the effective consortia not only significantly reduced the symptoms in treated leaves, but also conferred a systemic protection against the pathogen. All single strains, except for PIP6, and combinations provided a comparable level of inhibition in untreated leaves, suggesting that protection was mediated by the plant’s defense response and could therefore be transferred to distant parts of the plant. Compared with single strains, the effects of consortia showed a higher stability as reflected by the lower standard deviation even at the systemic scale, with the exception of the combination CHP2/CHP14/SOD2 (*Cupriavidus metallidurans, Bacillus subtilis, Bacillus halotolerans*) whose variability was comparable to strains used alone both at a local and systemic level, probably because of the negative impact of SOD2, as discussed above. While mechanisms involved in this systemic response triggered by consortia were not analyzed in the whole plant experiment, the data obtained from single strains suggest that they rely on the induction of the SAR pathway and on increased production of stilbenic compounds.

The use of combinations of bacteria with other bacterial strains, microorganisms from different kingdoms or even chemical compounds has been reported as a successful strategy against several plant pathogens in the past. This solution has several advantages, as it allows the addition of several effects such as direct pathogen inhibition, accumulation of more diverse antipathogenic compounds, plant defense stimulation and higher competition for space and resources (Niu et al., 2020). Strains of *Bacillus mycoides* and *Pichia guillermondii* were more effective at inhibiting *B. cinerea* spore germination and reducing necrotic lesion size in strawberry leaves when used in combination compared to their use alone, showing that organisms from different kingdoms can synergize and also provide more consistent protection (Guetsky et al., 2001). In this study, the level of protection conferred by the consortia often matched or exceeded the protection conferred by the best performing strain alone, and decreased the variability despite the dilution (OD_600_ = 0.3 for each strain in a consortium instead of OD_600_ = 1 for single strains), suggesting that strains assembled in a consortium have synergies and display better performance in terms of protection against diseases. From literature data, some genera seem to harbor more effective strains than others in terms of plant protection, as e.g. for bacteria, the genera *Bacillus* and *Pseudomonas* (Niu et al., 2020). The genus *Bacillus* has been reported as one of the four bacterial taxa whose presence was correlated with lower disease incidence severity in vineyard soils (Fournier et al., 2025). In our study, several different *Bacillus* species were proven effective but no *Pseudomonas* were present. Instead, other genera such as *Cupriavidus*, *Herbaspirillum*, *Microbacterium*, and members of other less commonly studied genera showed high protective potential. These results are consistent with the idea that optimal protection with bacteria-based biocontrol solutions may benefit from the use of combinations including diverse genera of beneficial microorganisms with complementary effects, and that strains showing only moderate effects when used alone could be of high importance to stabilize or increase the effectiveness of a consortium (Maciag et al., 2023; Nunes et al., 2024).

## Conclusion

Bacteria isolated from the grapevine phyllosphere represent a wide range of strains that, alone or in combination, significantly reduced symptoms caused by the two pathogens *P. viticola* and *B. cinerea* and were also able to interfere with the pathogens’ spore motility and germination. These strains were also able to stimulate their host defenses through the upregulation of defense-related genes and through the increased production of stilbenes.

The most effective bacteria consortia: CHP2/PID6/PIP6 (*C. metallidurans/B. cereus/H. huttiense*) and CHP14/PID13/PIP6 (*B. subtilis/B. endophyticus/H. huttiense*) caused a symptom reduction of 88 to 94 % *in planta* under growth chambers conditions, similar to that achieved with commercially available solutions used as a benchmark. This work suggests that the use of bacterial combinations is a valid strategy to increase biocontrol effectiveness. Such strategy combined with an adapted should be tested under field conditions to evaluate its suitability as a sustainable alternative to current crop protection practices.

## Acknowledgements

The authors are grateful to Alisson Gillon and Gaëlle Ruffieux for their help in the experimental work with grapevine infection and bacterial inoculum preparation.

